# Phasing in and out of phytophagy: phylogeny and evolution of the family Eurytomidae (Hymenoptera: Chalcidoidea) based on Ultraconserved Elements

**DOI:** 10.1101/2025.01.29.635542

**Authors:** Y. Miles Zhang, Gérard Delvare, Bonnie B. Blaimer, Astrid Cruaud, Jean-Yves Rasplus, Seán G. Brady, Michael W. Gates

**Affiliations:** Systematic Entomology Laboratory, USDA-ARS, c/o National Museum of Natural History, Washington DC, USA; Department of Entomology, Smithsonian Institution, National Museum of Natural History, Washington, DC, USA; UMR CBGP, CIRAD, F-34398, Montpellier, France; CBGP, CIRAD, INRAe, IRD, Montpellier SupAgro, Université de Montpellier, Montpellier, France; Museum für Naturkunde, Leibniz Institute for Evolution and Biodiversity Science, Center for Integrative Biodiversity Discovery, Berlin, Germany

**Keywords:** UCE, entomophytophagy, seed chalcid, dating, parasitoids

## Abstract

We present the first global molecular phylogenetic hypothesis for the family Eurytomidae, a group of chalcidoid wasps with diverse biology, with a representative sampling (197 ingroups and 11 outgroups) that covers all described subfamilies, and 70% of the known genera. Analyses of 962 Ultra-Conserved Elements (UCEs) with concatenation (IQ-TREE) and multispecies coalescent approaches (ASTRAL) resulted in highly supported topologies in recovering the monophyly of Eurytomidae and its four subfamilies. The taxonomy of Eurytomidae, and in particular the large subfamily Eurytominae, needs major revisions as most large genera are recovered as para- or polyphyletic, and the erection of multiple new genera is required in the future to accommodate these taxa. Here, we synonymize the genera *Cathilaria* (*C. certa, C. globiventris, C. opuntiae*, and *C. rigidae*) and the monotypic *Aiolomorphus rhopaloides* within *Tetramesa* **syn. nov**., *Parabruchophagus* (*P. kazakhstanicus, P. nikolskaji, P. rasnitsyni, P. saxatilis*, and *P. tauricus*) and *Exeurytoma* (*E. anatolica, E. caraganae*, and *E. kebanensis*) within *Bruchophagus* **syn. nov**.. We also provide 137 DNA barcode *COI* fragments extracted from the UCE contigs to aid in future identifications of Eurytomidae using this popular genetic marker. Eurytomidae most likely originated in South America with an estimated crown age of 83.37 Ma. Ancestral state reconstruction indicates that secondary phytophagy has evolved at least seven times within the subfamily Eurytominae, showcasing the evolutionary flexibility of these vastly understudied wasps.

## Introduction

The family Eurytomidae encompasses more than 1600 described species distributed across 77 genera, organized into four subfamilies (Gates et al., 2025; UCD Community, 2025). Among these subfamilies, Eurytominae, the largest one, comprises 66 genera and approximately 1500 species (UCD Community, 2025). In contrast, the remaining three subfamilies, Heimbrinae (two genera/eight species) (Stage & Snelling, 1986), Rileyinae (seven genera/78 species) (Gates, 2008), and Buresiinae (two genera/nine species) (Zerova, 1988), are considerably smaller. Eurytomidae are identified by their quadrate pronotum, head and mesosoma with umbilicate sculpture, mesothoracic spiracle not exposed, and generally black, brown, or yellow coloration. For a more in-depth examination of morphology, refer to Gates et al. (2025). However, identification of eurytomid genera is often difficult due to the seemingly uniform appearance of the species and the blending of character traits that often form morphoclines. The largest genus *Eurytoma* Illiger includes over half of the described eurytomid species, while more than 40 genera are monotypic (Lotfalizadeh et al., 2007). To make matters worse, molecular data is lacking for most eurytomid genera in public databases, making it difficult for non-experts to identify specimens accurately. This is partly due to the low success rate in amplifying the typical barcoding fragment using universal primers (e.g., Zhang et al., 2014), although recently, primer sets with higher success rates have been designed specifically for eurytomids (Zhang et al., 2022; Jafari et al., 2023).

Eurytomids are present on every continent except Antarctica, with the highest diversity observed in tropical regions, followed by temperate climates. They showcase a wide range of biological characteristics and exhibit rapid evolution in both host utilization and feeding behavior. The majority of eurytomid larvae are endophytic, serving as seed eaters, gall formers, or parasitoids of phytophagous insects (Gates et al., 2025). Most species act as primary or secondary parasitoids, targeting eggs, larvae, or pupae of various arthropod groups such as Diptera, Coleoptera, Hymenoptera, Lepidoptera, Orthoptera, and Araneae. Host-parasitoid relationships for many endophytic species remain unclear, and some Eurytomidae believed to be parasitoids may function as inquilines (Lotfalizadeh et al., 2007). A subset of eurytomid species holds economic significance, either as potential biological control agents or pests. Due to their diverse biology, Eurytomidae play contrasting roles in agriculture. Many are harmful seed-eaters affecting cultivated plants, while others are used for the biological control of invasive plants (Gates et al., 2025).

The monophyly of Eurytomidae and the relationships between the subfamilies remain debated as different morphological and molecular data have provided conflicting answers (Gates, 2008; Lotfalizadeh et al., 2007). The phylogenetic analysis of Gates (2007) used 50 morphological characters with a sampling focus on Rileyinae, recovering a monophyletic family. Lotfalizadeh et al. (2007) used 150 morphological characters and focused on the large subfamily Eurytominae, instead recovering a paraphyletic Eurytomidae, with Heimbrinae as the sister to Chalcididae, another family of chalcid wasps used as outgroup, instead of grouping with the rest of the family. Using multilocus data, Campbell et al. (2000) included five eurytomid samples within their Chalcidoidea phylogeny inferred using the *28S-D2* rDNA, and recovered a polyphyletic Eurytomidae comprising of two disparate groups, Rileyinae and Eurytominae. Chen et al. (2004) published the first molecular phylogeny of the family using 24 species and four genes, *18S* and *28S* (ribosomal DNA) and *16S* and *COI* (mitochondrial DNA). While most genes exhibited very low levels of variability and the ribosomal DNA results strongly conflicted, the authors concluded that the family was not monophyletic, with Rileyinae sister to Dirhininae (Chalcididae) and nested within a clade containing other chalcidoid families such as Eunotidae, Perilampidae, and Eucharitidae. Munro et al. (2011) also failed to recover a monophyletic Eurytomidae in their molecular analysis of Chalcidoidea using *18S* and *28S*, as while Rileyinae was recovered as a monophyletic group, Buresiinae and Heimbrinae did not cluster with the other Eurytomidae. Heraty et al. (2013) recovered a monophyletic Eurytomidae using a combination of morphology with the same two ribosomal genes, with all subfamilies monophyletic except for Heimbrinae which placed internally with *Neorileya* Ashmead (Rileyinae). Using transcriptome data, Peters et al. (2018) recovered a monophyletic Eurytomidae as sister to Chalcididae using transcriptomic data and a reduced representation matrix, but their sampling included only two specimens/genera of Eurytominae.

With the rapid advances in sequencing technology, it is now possible to generate thousands of loci from non-destructively sampled museum specimens (Cruaud et al., 2019), enabling the increasing popularity of museum-based phylogenomic studies dubbed “museomics”. Target capture approaches such as Ultraconserved Elements (UCEs, Faircloth et al., 2012) have revolutionized our understanding of arthropod phylogenomics, including studies on Hymenoptera and/or Chalcidoidea that have included Eurytomidae (e.g., Blaimer et al., 2023; Branstetter, Longino, et al., 2017; Cruaud et al., 2021; Cruaud et al., 2024; Zhang et al., 2020; Zhang et al., 2022). However, the monophyly of Eurytomidae remained unresolved in these studies, until Cruaud et al. (2024) recovered a monophyletic Eurytomidae (18 taxa representing all four subfamilies) with Chalcididae as sister-group. Conversely, the Blaimer et al. (2023) study used many of the same samples as Cruaud et al. (2024) but did not recover Eurytomidae as monophyletic despite a wider sampling of 35 taxa; *Heimbra opaca* (Ashmead) behaved as a rogue taxon in this study by grouping either with the colotrechnine pteromalid *Colotrechnus* Thomson or within Chalcididae instead of within the otherwise monophyletic Eurytomidae.

The goal of the current study is to use UCEs and an expanded global taxonomic sampling in order to 1) test the monophyly of Eurytomidae, the four subfamilies, and major genera; 2) infer divergence times of major groups using molecular dating analyses, and 3) to investigate how phytophagy and entomophagy have evolved over the history of the family.

## Materials & Methods

### Taxonomic Sampling

The dataset comprises 208 individuals with 197 ingroups chosen to cover the taxonomic breadth within Eurytomidae. UCE data were newly generated for 151 ingroups while data for 57 specimens were acquired from previously published studies, including 10 Chalcididae and one Chalcedectidae as outgroups (Blaimer et al., 2023; Branstetter, Danforth, et al., 2017; Cruaud et al., 2021; Cruaud et al., 2024; Zhang et al., 2022). The ingroup taxa represents 48 genera, as well as a number of specimens which could not be placed to genus within the current classification (Table S1, Supplemental File 1). The identification of the specimens were confirmed by authors considered as the world authorities on Eurytomidae (GD and MWG). Twenty-one known genera are missing from our sampling, but these genera constitute only 49 species among the >1600 total described species from the family (Supplemental File 1). In many cases these missing genera are monotypic and known only from their type specimens. A few other missing genera are doubtful according to the original descriptions. Nevertheless some regions are under-sampled, especially the tropical realms (Neotropical, Afrotropical and Indomalayan regions) mostly because of the lack of suitable material for the sequencing.

### Generation of the UCE Dataset

The UCE library preparation (N=151) was conducted in the Laboratories of Analytical Biology at the Smithsonian Institution’s National Museum of Natural History (NMNH, Washington DC, USA) and Centre de Biologie pour la Gestion des Populations (GBCP, Montpellier, France). The protocol largely follows the standard pipeline for capturing and enriching UCE loci from Hymenoptera (Branstetter, Longino, et al., 2017; Cruaud et al., 2019; Zhang et al., 2019). Briefly, DNA was extracted using the DNeasy Blood and Tissue Kit, with individuals either destructively processed during extraction or left intact. The Kapa Hyper Prep library preparation kit (Kapa Biosystems, Wilmington, MA) was used along with TruSeq universal adapter stubs and 8-bp dual indexes (Glenn et al., 2019), combined with sheared genomic DNA and amplified using polymerase chain reaction (PCR). We followed the myBaits V4 or V5 protocol (Arbor Biosciences, Ann Arbor, MI) for target enrichment of the pooled DNA libraries (4–12 samples per pool) using either the Hymenoptera 1.5Kv1 (Faircloth et al., 2015) or the 2.5Kv2P kit (Branstetter, Longino, et al., 2017), with probe/target hybridization at 65°C for 24 h. The combined library was sequenced on an Illumina HiSeq 2500 (150-bp paired-end) at Huntsman Cancer Institute (Salt Lake City, UT, USA) or Admera Health (South Plainfield, NJ, USA); or on an Illumina MiSeq (300-bp paired-end) at UMR AGAP (Montpellier, France). Voucher specimens are stored either at NMNH, GBCP, the Centre de Coopération International en Recherche Agronomique pour le Développement (CIRAD, Montepellier, France), or Taiwan Agricultural Research Institute (TARI, Taichung, Taiwan).

### UCE Processing and Matrix Assembly

All downstream analyses were conducted on the Smithsonian High Performance Cluster (SI/HPC, https://doi.org/10.25572/SIHPC). We used the Phyluce v 1.7.4 pipeline (Faircloth, 2015) to process newly generated raw reads and published genome/UCE assemblies mined from previous studies. Adapters were trimmed using illumiprocessor or trimmomatic (Bolger et al., 2014; Faircloth, 2013), and assembled using spades v 3.14.0 (Prjibelski et al., 2020). The assemblies were aligned using MAFFT v 7.490 (Katoh et al., 2002; Katoh & Toh, 2008), and trimmed using GBlocks (Castresana, 2000) using the following settings: b1 = 0.5, b2 = 0.5, b3 = 12, b4 = 7. Additionally, we used Spruceup v 2024.7.22 with 99% Weibull distribution or manual cutoff of select samples to remove any potentially misaligned regions as they can produce exaggerated branch lengths (Borowiec, 2019a). The manual cutoffs samples and their values are: (*Aximopsis*_sp1_USNMENT01322402, 0.4; *Aximopsis*_sp2_USNMENT01322403, 0.4; *Bruchodape*_sp_USNMENT01322364, 0.4;

*Bruchophagus*_*abscedus*_USNMENT01525852, 0.35;

*Bruchophagus*_*gibbus*_USNMENT01322388, 0.25;

*Bruchophagus*_*phlei*_USNMENT01322376, 0.4; *Bruchophagus*_*roddi*_USNMENT01322386, 0.3;

*Burksoma*_*scimitar*_USNMENT01322361, 0.5; *Eudoxinna*_sp_USNMENT01322401, 0.4;

*Eurytoma*_*erythrinae*_USNMENT01938343, 0.4;

*Eurytoma*_*obtusiventris*_USNMENT01322387, 0.4; *Heimbra*_*opaca*_USNMENT01339599, 0.6;

New_Genus_USNMENT01322400, 0.5; *Philolema*_*latrodecti*_USNMENT01322374, 0.4; and *Phylloxeroxenus*_sp3_USNMENT01322411, 0.5). Additionally, DNA barcode sequences were bioinformatically extracted from UCE contigs using phyluce_assembly_match_contigs_to_barcodes.

### Phylogenetic Analyses

We selected the 50% complete matrix (1605 loci) and 70% complete matrix (962 loci) which represents loci that are present in >50% and _≥_70% of the taxa, respectively. The 70% matrix is our preferred dataset and is used for other downstream analyses, and while the 50% consisted of a larger number of genes, it also had higher levels of missing data (for results of all 50% analyses see Figures S5–8). Phylogenetic summary statistics were calculated using AMAS v 0.98 (Borowiec, 2016). We conducted phylogenetic analyses under the maximum-likelihood criterion with IQ-TREE v 2.3.1 with the default 100 likelihood searches (Minh et al., 2020a). The best model for each loci were selected using ModelFinder v2.0 (Kalyaanamoorthy et al., 2017) using the IQ-TREE command “-MFP+MERGE”. To assess support, we performed 1000 Ultrafast Bootstrap (UFBoot, Hoang et al., 2017), along with “-bnni” to reduce the risk of overestimation, and a Shimodaira–Hasegawa approximate likelihood-rate test (SH-aLRT, Guindon et al., 2010) with 1000 replicates. Only nodes with support values of UFB _≥_95 and SH-aLRT _≥_80 were considered robust. We also inferred gene and site concordance factors (gCF and sCF) following Minh et al (2020b) resulting in the trees 50p_CONCORD and 70p_CONCORD.

In order to minimize the effects of phylogenetic heterogeneity, we also filtered the matrices with the symmetry test of stationarity, reversibility, and homogeneity for sequence alignments (Naser-Khdour et al., 2019), and conducted separate analyses using the same settings as above resulting in the trees 50p_SYM and 70p_SYM. The matrices were also analyzed using partitions based on Sliding-Window Site Characteristics of Site Entropy (SWSC-EN, Tagliacollo & Lanfear, 2018), and partitioned using the rcluster algorithm in PartitionFinder2 via RAxML using default settings (Lanfear et al., 2014; Lanfear et al., 2016; Stamatakis, 2006) resulting in the trees 50p_SWSC and 70p_SWSC. We also extracted the protein-coding genes using the custom scripts provided by Borowiec (2019b) using the protein FASTA file from assembled genomes of *Nasonia vitripennis* (Walker) as reference. Analysis of the resulting concatenated matrix of amino acids using the posterior mean site frequency (PMSF) method in order to ameliorate long-branch attraction artefacts (Wang et al., 2018). The 50p_SWSC and 70p_SWSC were used as guide trees with 60-component mixture models (Wang et al., 2018), resulting in trees 50p_PMSF_C60 and 70p_PMSF_C60, respectively.

Species trees under the multi-species coalescent (MSC) model were inferred using ASTRAL-III v5.7.8 (Zhang et al., 2018) for both matrices. Gene trees of each locus were estimated in IQ-TREE, and the best models of nucleotide substitution were selected using ModelFinder. We collapsed all branches in the gene trees with <10% UFB using the ‘nw_ed’ command included in Newick Utilities (Junier & Zdobnov, 2010), as this step has been shown to improve accuracy in MSC analysis (Zhang et al., 2018). Support was assessed by annotating the MSC tree with local posterior probabilities (LPP, Sayyari & Mirarab, 2016), with _≥_0.95 considered as strong support, resulting in trees 50p_BS10_ASTRAL 70p_BS10_ASTRAL.

Resulting raw tree outputs were modified using R packages ggtree (Yu et al., 2017), treeio (Wang et al., 2020), and phytools (Revell, 2024), along with Inkscape v1.4.

### Divergence Dating

We estimated the divergence times using MCMCTree using PAML v4.9j (Yang, 2007), using approximate likelihood calculation and ML estimation of branch lengths and the 70p_SWSC tree as the fixed input along with the 70% matrix. Due to the lack of reliable published eurytomid fossil records, we instead used secondary calibration based on divergence dates from previous studies (Blaimer et al., 2023; Cruaud et al., 2024). The soft minimum bound for the common ancestor of Eurytomidae + Chalcididae was set at 92Ma, with Chalcididae and Eurytomidae set at 73–86 Ma and 74–84 Ma, respectively (Blaimer et al., 2023; Cruaud et al., 2024). Uniform distributions were used as calibration densities. We conducted the analysis with a 50,000-generation burnin and 100,000 additional generations using independent rates under the HKY model to strike balance between complexity and computational efficiency, and convergence was assessed using Tracer v1.7.1 (Rambaut et al., 2018) to ensure ESS values were >200 for all categories. The resulting tree was visualized using MCMCtreeR (Puttick, 2019) in R v4.3.3 (R Core Team, 2024).

### Historical Biogeography

Historical biogeography was estimated using BioGeoBEARS v1.1.1 (Matzke, 2014; 2018) using the MCMCTree result as input. Coding of occurrences were based on collecting labels and expert knowledge, and scored as present/absent in six biogeographic regions (Neotropical, Nearctic, Afrotropical, Palearctic, Indomalayan, and Australasian) (Table S3). Dispersal-Extinction-Cladogenesis (DEC; Ree and Smith, 2008), BAYAREALIKE, and DIVALIKE (Ronquist, 1997) models were used with and without the jump parameter for founder events (+J; Matzke, 2014), and maximum range size allowed were set to five. Model selection was performed based on AICc (Matzke, 2022). We included constraints on three different time periods (0-23 Ma/Neogene + Quaternary; 66–23 Ma/Paleogene; and 100–66 Ma/Late Cretaceous) with different dispersal rate scalers following Cruaud et al. (2024) (Table S5a).

### Ancestral State Reconstruction of Feeding Behavior

We reconstructed the ancestral states of feeding behavior using the R package CorHMM v2.8 (Beaulieu et al., 2013), using the 70p_SWSC phylogeny as input after pruning the outgroups. We coded “0” for entomophagous, “1” for phytophagous, and “?” for unknown (Table S3). The host data was based on a combination of sample information, literature, and unpublished data (Table S3). We performed reconstructions under the “equal rates” model (ER), “symmetrical rate” (SYM) and “all rates different” model (ARD) and compared the fit of these models with the corrected Akaike Information Criterion (AICc).

## Results & Discussion

### Phylogenetic Relationships of Eurytomidae

Our study is the most comprehensively sampled phylogenomic study of Eurytomidae to date, highlighting the utility of museomics on poorly studied but diverse lineages of small parasitic wasps. The number of UCE loci recovered ranged from 228–2145 (Table S1). The 50% complete UCE matrix consisted of 565,920bp of data, of which 414,715 (73.3%) are variable sites, and 324,331 (57.3%) are parsimony informative. The 70% complete UCE matrix consisted of 340,005bp of data, of which 255,163 (75%) are variable sites, and 204,108 (60%) are parsimony informative. The amount of missing data for the 50% and 70% matrices was 38.82% and 32.04%, respectively (Table S2). We also managed to extract 137 *COI* fragments >200 bp from the UCE assemblies, and while these fragments cannot be used to accurately infer family level relationships, they can still improve the publicly available sequences of Eurytomidae for future identifications.

The monophyly of Eurytomidae and its four subfamilies was recovered in all of our analyses (Figure 1, S1–8), which is consistent with Cruaud et al. (2024) and somewhat consistent with previous morphological studies (Gates 2007; Lotfalizadeh et al., 2007). The topologies of the 50% and 70% matrices are mostly identical among the different analyses, with slight shifts among the relationships among genera within the *Chryseida* clade of eurytomine (Figure 1, Figure S1,S5). Larger differences were observed between the protein-coding-only analyses (Figure S3,S7); and between the MSC analyses using ASTRAL (Figure S4,S8). Here we summarize our main findings based on the preferred tree of the 70% matrix partitioned by SWSC (70p_SWSC) as it limits the impact of missing data and model violations, while also highlighting major differences among analyses (for full comparisons see Figure S1–8):

**Figure 1.**
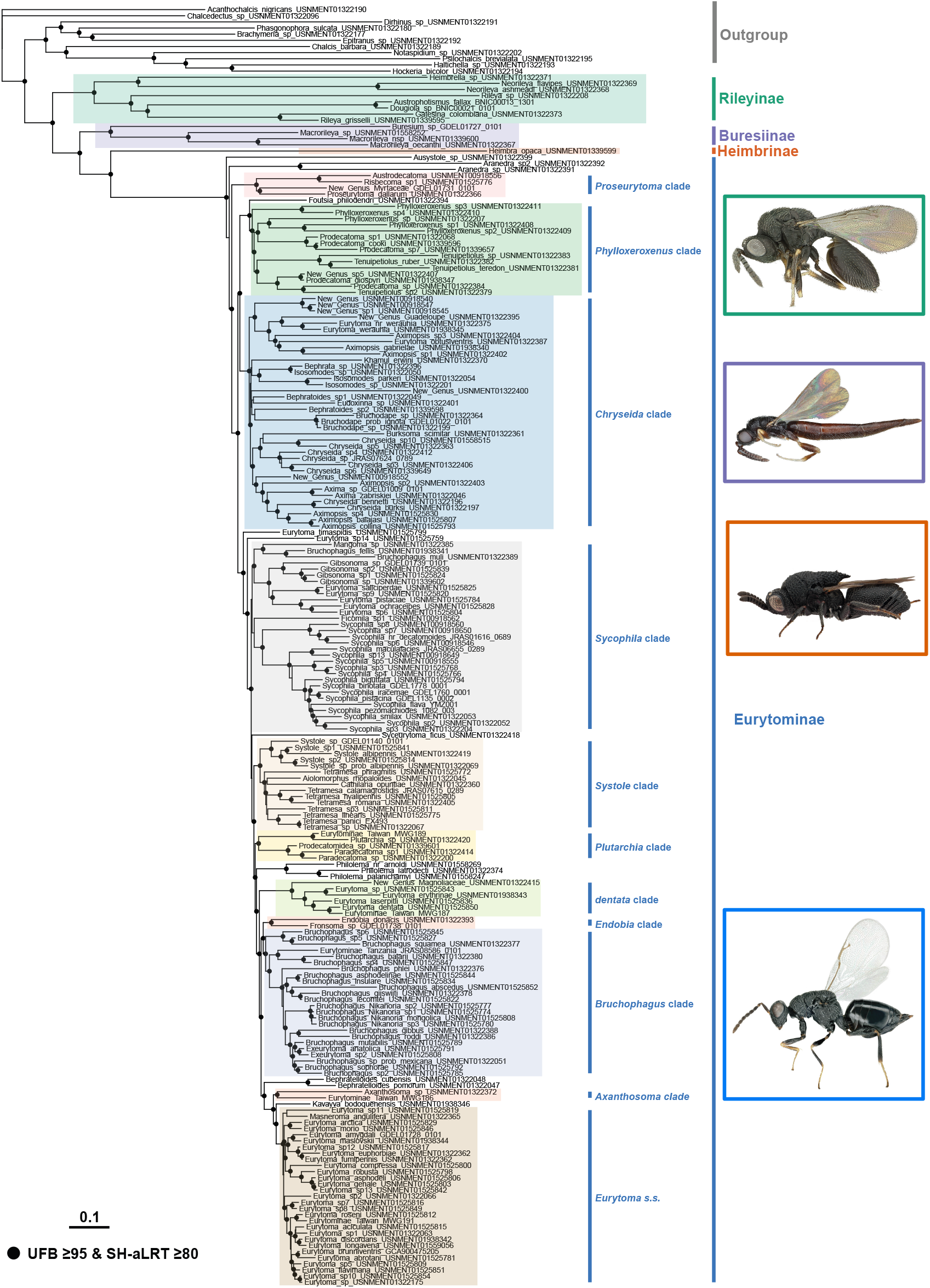
Phylogram of Eurytomidae based on the 70% UCE matrix, inferred under the SWSC-EN partitioning, partition merging, and ModelFinder model selection, generated using IQ-TREE (70p_SWSC). Black dots at nodes indicate support values of UFB _≥_95 and SH-aLRT _≥_80, values below that are omitted. Scale bar in substitutions per site. Habitus photos from top to bottom: *Rileya pallidipes* (Miles Zhang), *Macrorileya* sp. (Miles Zhang), *Heimbra opaca* (Craig Brabant), and *Eurytoma abrotani* (Gérard Delvare).

Rileyinae is recovered to be the sister to all other eurytomids. Among the two genera within this subfamily represented by more than one species, *Neorileya* was monophyletic while *Rileya* Ashmead was not.

Buresiinae was recovered as the sister group to Heimbrinae and Eurytominae, and within this subfamily *Macrorileya* Ashmead was recovered as a monophyletic group in the ML analyses and sister to *Buresium* Bou_č_ek, which was represented by a single specimen. *Macrorileya_*sp_USNMENT01558252 is recovered as the sister group to the rest of eurytomine in the MSC analysis with strong support of LPP=1, as opposed to being grouped with the rest of *Macrorileya*.

Heimbrinae was only represented by *H. opaca* in our study, but it was recovered as the sister to Eurytominae in all analyses except in the 70p_PMSF_C60 tree, where it was recovered as the sister to Rileyinae. We used the same sample of *Heimbra_opaca*_USNMENT01139599 as in Blaimer et al. (2023) as the sole representative for Heimbrinae. While it was recovered outside of Eurytomidae in that earlier study; it was placed as the sister to the subfamily Eurytominae in all of our analyses similar to Cruaud et al. (2024). The difference in topology with the Blaimer et al (2023) could be the result of filtering of misaligned regions, or better taxonomic sampling.

The subfamily Eurytominae is the largest of the four subfamilies and represents the bulk of our taxonomic sampling. Unsurprisingly, most of the larger genera where we included multiple species were not recovered as monophyletic: *Aximopsis* Ashmead, *Bruchophagus* Ashmead, *Chryseida* Spinola, *Eurytoma, Gibsonoma* Narendan, *Isosomodes* Ashmead, *Prodecatoma* Ashmead, *Phylloxeroxenus* Ashmead, and *Tetramesa* Walker. *Bephratelloides* Girault, *Bephratoides* Brues, *Bruchodape* Burks, *Exeurytoma* Burks, *Paradecatoma* Masi, *Philolema sensu largo* Cameron, *Sycophila* Walker, and *Systole* Walker were monophyletic. The relationships among genera within the large subfamily Eurytominae is somewhat consistent with the morphological study by Lotfalizadeh et al. (2007); we provide more detailed taxonomic discussion in the Supplemental File. This study highlights the need to revise the generic concepts for Eurytominae, as many genera are para- or polyphyletic. While this is a well-known problem within the subfamily, the high level of morphological homoplasy and interpretation of difficult morphological characters made it extremely challenging to accomplish (Lotfalizadeh et al., 2007). With the aid of the UCE phylogeny as a backbone, refinement of Eurytominae taxonomy can take place on a smaller scale with more extensive taxonomic sampling of certain clades of interest (Zhang et al., 2021). The establishment of multiple new genera are needed to accommodate outliers that do not fit the current generic limits (e.g., Campos-Moreno et al., 2022; Gates & Cascante-Marin, 2004; Gates & Delvare, 2008), preferably in combination with UCE and/or multilocus data such as *COI*.

The base of the Eurytominae consists of a series of phytophagous genera: 1) The genus *Ausystole* Bouček is sister to all other Eurytominae, 2) followed by the genus *Aranedra*, 3) then the *Proseurytoma* clade consisting of *Austrodecatoma* Girault, *Risbecoma* Subba Rao, *Proseurytoma* Kieffer, and an undescribed genus feeding on Myrtaceae.

The next clade consists of the genus *Foutsia* Burks, then a paraphyletic assemblage of a clade that includes *Phylloxeroxenus, Prodecatoma*, and *Tenuipetiolus* Bugbee. This is followed by the large *Chryseida* clade that consists of a mix of some ‘*Aximopsis’* (including the newly described microgastrine braconid hyperparasitoid ‘*Aximopsis’ gabrielae* Zhang, Gates & Campos) and ‘*Eurytoma’* species (parasitoid ‘*Eurytoma’ obtusiventris* Gahan, phytophagous ‘*Eurytoma’ werauhia* Gates, and New_Genus_Guadeloupe_USNMENT01322395), then *Eudoxinna* Walker and an undescribed genus assigned to the ‘*erythroapsis* group’ New_Genus_USNMENT01322400. This is followed by ((*Bephrata* Cameron *+ Isosomodes*) *+ Khamul* Gates), which are sister to (*Bephratoides + Bruchodape*), and all of which are sister to a clade with *Axima* Walker, the remaining *Aximopsis, Chryseida*, and the monotypic *Burksoma scimitar* Subba Rao. In the PMSF analysis, *Bephratoides_*sp2_USNMENT01339598 was instead recovered as the sister to *Eudoxinna*, while *Khamul* was an isolated branch.

*‘Eurytoma’ timaspidis* (Mayr) (*aspila* species group) and *Eurytoma_*sp14_USNMENT01525759 (*verticillata* species group) were recovered as isolated branches, and are broadly separated from the bulk of *Eurytoma* including its type species *Eurytoma abrotani* (Panzer). This is followed by the *Sycophila* clade, which consists of (‘*Bruchophagus’* feeding on *Citrus* + *Mangoma* Subba Rao) + (‘*Eurytoma’* in the *salicis* and *pistaciae* species group + *Gibsonoma*); are sister to (*Sycophila + Ficomila* Bouček).

The monotypic *Syceurytoma ficus* Bouček is recovered as an isolated lineage, followed by the *Systole* clade which consists of the phytophagous *Systole*, a monophyletic genus. Its sister group *Tetramesa*, is largely monophyletic with the exception of *Tetramesa phragmitis* (Erdös), which is sister to all of other *Tetramesa* plus the monotypic *Aiolomorphus rhopaloides* Walker and *Cathilaria opuntiae* (Muesebeck). As all these genera gall twigs or flowers of Poaceae, the four described *Cathilaria* species (*C. certa* Zerova, *C. globiventris* (Zerova), *C. opuntiae*, and *C. rigidae* Zerova) and *A. rhopaloides* are synonymized within *Tetramesa* **syn. nov**., which is consistent with their biology and no longer renders *Tetramesa* paraphyletic.

The *Plutarchia* clade includes the genus *Plutarchia* Girault, unplaced Eurytominae_Taiwan_MWG189, *Prodecatomidea* Risbec and two species of *Paradecatoma*, followed by the clade/genus *Philolema s*.*l*..

The ‘*Eurytoma’ dentata* clade consists of an unnamed genus reared from *Magnolia* L., *Eurytoma erythrinae* Gates & Delvare, *Eurytoma laserpitii* Mayr, *Eurytoma dentata* Mayr, and unplaced Eurytominae_Taiwan_MWG187.

The small *Endobia* clade (*Fronsoma* Narendran + *Endobia* Erdös) is sister to the *Bruchophagus* clade, which consists of the rest of *Bruchophagus*, unplaced

Eurytominae_Tanzania_JRAS08586_0101, and *Exeurytoma*. In the 70p_PMSF_C60 tree, *Bruchophagus roddi* (Gussakovskiy) was recovered as an isolated lineage outside of these two clades, whereas in the 70p_PMSF_C60 tree, *Fronsoma* was recovered in this position. The *Bruchophagus bajarii, gibbus, phlei*, and *squamea* species groups defined by Lotfalizadeh et al. (2007), along with the former genus *Nikanoria* (Nikol’skaya) were recovered within *Bruchophagus*. Additionally, a set of *Bruchophagus* species reared from seeds of Asphodelaceae sharing the same characteristics as the genus *Parabruchophagus* Zerova (*B. lecomtei* Delvare, *B. gijswijti* Askew & Ribes, *B. abscedus* Askew, *B. insulare* Delvare, *B. asphodelinae* Askew & Stojanova) is recovered within the genus. Therefore, all five described species included in *Parabruchophagus* (*P. kazakhstanicus* Zerova, *P. nikolskaji* Zerova, *P. rasnitsyni* Zerova, *P. saxatilis* Zerova, and *P. tauricus* Zerova), which also develop in the seeds of Asphodelaceae (*Eremurus* M.Bieb.), are transferred to *Bruchophagus* **syn. nov**.. Finally, *Exeurytoma* which is represented by *E. anatolica* Cam and an undescribed species *Exeurytoma_*sp2_USNMENT01525808 are recovered as the sister to the *Bruchophagus gibbus* species group, all three described *Exeurytoma* species (*E. anatolica, E. caraganae* Burks, and *E. kebanensis* Doganlar) are therefore synonymized within *Bruchophagus* **syn. nov**. as both groups develop within leguminous pods.

The final grouping consists of the clade/genus *Bephratelloides* Girault, followed by *Axanthosoma* clade (unplaced Eurytominae_Taiwan_MWG191 and *Axanthosoma* Girault), the newly described *Kavayva* Zhang, Silvestre & Gates, and finally the rest of *Eurytoma sensu stricto* (including unplaced Eurytominae_Taiwan_MWG191) + the monotypic *Masneroma angulifera* Bouček. In the 70p_PMSF_C60 tree, *Eurytoma_*sp11_USNMENT01525819 was instead recovered outside of *Kavayva*/*Eurytoma sensu stricto*/*Masneroma* clade. The following *Eurytoma s*.*s*. species groups from Lotfalizadeh et al. (2007) were recovered within our analysis: *robusta, compressa, amygdali, morio, fumipennis, rosae/arbortani* + *appendigaster* groups.

### Divergence Dating & Historical Biogeography

Based on our divergence dating analysis (Figure 2, S9, Table S4), the crown age of Eurytomidae is ∼83.3 Ma [95% equal tail credibility interval (CI) 81.1–84.9 Ma], with Rileyinae estimated at ∼71.8 Ma (64.9–80.0 Ma); followed by Buresiinae at ∼66.6 Ma (55.6–75.6 Ma). The subfamilies Eurytominae and Heimbrinae diverged at ∼75.5 Ma (71.5–80.2 Ma), while crown Eurytominae diverged at ∼70.9 Ma (66.6–75.8 Ma).

**Figure 2.**
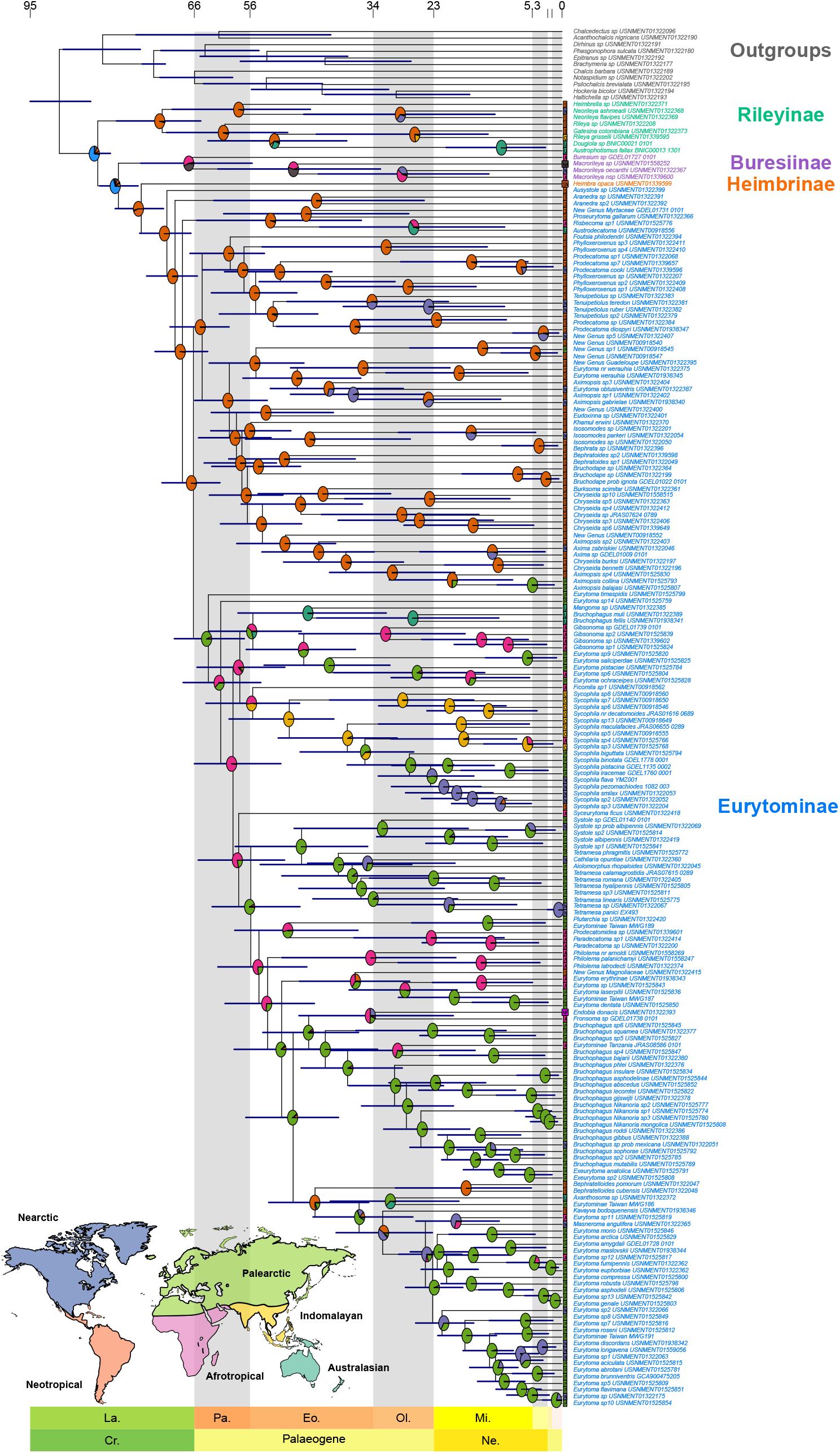
Chronogram of Eurytomidae inferred in MCMCTree under likelihood approximation, root age with hard bound at 91.5 Ma, and independent rates clock using fixed consensus topology of 70p_SWSC. Age range in 95% Highest Posterior Density (HPD) is shown as blue bars. Ancestral biogeographic reconstruction using the DIVALIKE+J model, with the geographic regions as Neotropical (S), Nearctic (N), Afrotropical (A), Palearctic (P), Indomalayan (O), and Australasian (U). Pie charts at nodes indicate likely geographic region assignments.

DIVALIKE+J was selected as the best model, with a weighted AICc of 0.48 (Table S5b). The historical biogeographic analyses recovered a South American origin for Eurytomidae (Figure 2, S10). This result aligns well with the paleogeographic configurations during the Late Cretaceous and early Cenozoic, where the three continents were still relatively connected through intermittent land bridges or close continental proximities. Rileyinae, Heimbrinae, and Eurytominae are proposed as having a South American origin, while Buresiinae was recovered with an Afrotropical origin. A major divide within Eurytominae occurred around 66 Ma around the K-Pg mass extinction, splitting the group into mostly Old World and New World clades. The timing of this divergence coincides with the rapid radiation of angiosperms, which began earlier in the Late Cretaceous (∼100–70 Ma) but saw major ecological restructuring after the K-Pg extinction event. Angiosperms provided new ecological opportunities and likely contributed to the diversification of Eurytominae, which are associated with plant-host interactions, particularly seeds and gall-forming species (Peris & Condamine 2024). This angiosperm-driven expansion is consistent with the Eurytomidae diversification patterns in both the Old and New World.

As the divergence dates of Eurytomidae were secondarily derived from previous studies (Blaimer et al., 2023; Cruaud et al., 2024), it is not surprising that the dates recovered in our study were similar to previous studies, estimating an age of ∼83 Ma for crown-group Eurytomidae. However, the ages of the split between Heimbrinae and Eurytominae and the crown age of Eurytominae are estimated to be much older in our study (∼75.5 Ma, and ∼70.9 Ma, respectively), when compared to Cruaud et al. (2024) (∼66.3 Ma and ∼33.5 Ma, respectively). This could be the result of more comprehensive taxonomic sampling in our study, or the lack of fossil calibrations within Eurytominae. Nevertheless, these results align with the biogeographic history of the Late Cretaceous and early Paleogene, during which South America and Africa were in the final stages of separating (∼100–80 Ma), and the Atlantic Ocean was widening. The mostly Neotropical origin of Eurytomidae is also consistent with Cruaud et al. (2024). In the Paleogene Eurytominae split into two major clades, one predominantly found in Americas (Nearctic + Neotropical), while the other has moved into the Old World (Palearctic, Afrotropical, Indomalayan, Australasian) with intermittent shifts back into the Nearctic. This transcontinental expansion likely reflects a combination of tectonic events, the development of tropical forests in the Paleogene, and the global dispersal of angiosperms, which facilitated the spread of plant-associated species (Benton et al. 2022; Peris & Condamine 2024). The role of plate tectonics in facilitating these shifts—such as the northward drift of India and the eventual collision with Asia—may have contributed to the diversification and establishment of the group in the Indomalayan and Australasian regions.

It is worth pointing out that while the Mid-Jurassic (169–162 Ma) is a common estimated age of origin for Chalcidoidea (Blaimer et al., 2023; Cruaud et al., 2024), there is a 30 Ma gap between this age and the oldest known chalcidoid fossil from the Early Cretaceous (∼128 Ma), thus more detailed studies are needed to confirm the validity of the divergence dates (Zhang et al., 2025; Rasplus et al., 2025). Eurytomidae is very rare in the fossil records, and to date the oldest confirmed fossil is an undescribed species from the French Oise fossil (55.8 – 48.6 Ma) (Brasero & Martin 2009), which likely represents an extinct lineage as it does not match any of the four extant subfamilies. Additionally, three more confirmed Eurytomidae fossils have been described: one from the Green River formation in Wyoming, USA (50.3 – 46.2 Ma), and two from the Florissant Shale in Colorado, USA (37.2 – 33.9 Ma) (Rasplus et al., 2025). All three of these species have been assigned to *Eurytoma*, but this will need to be reexamined given the polyphyletic nature of the genus. Nevertheless, the rapid diversification of angiosperms and other phytophagous insects during the Cretaceous, coupled with the continued breakup of Pangea, likely enabled Eurytomidae to achieve a cosmopolitan distribution and develop such diverse host ranges, warranting further study.

### Evolution of secondary phytophagy

We tested three different models of ancestral state reconstruction, with the ARD model receiving the lowest AICc score (Figure 3, Table S6). The ancestral state of Eurytomidae is entomophagy, which is observed in Rileyinae, Buresiinae, and Heimbrinae. Phytophagy is likely the ancestral state of the subfamily Eurytominae as the earliest branching lineages (e.g., *Ausystole, Aranedra, Proseurytoma*) are all phytophagous. Entomophagy has evolved at least seven times within Eurytominae, including genera such as *Aximopsis, Chryseida, Eurytoma s*.*s*., *Philolema s*.*l*., *Sycophila*, and *Tenuipetiolus*.

**Figure 3.**
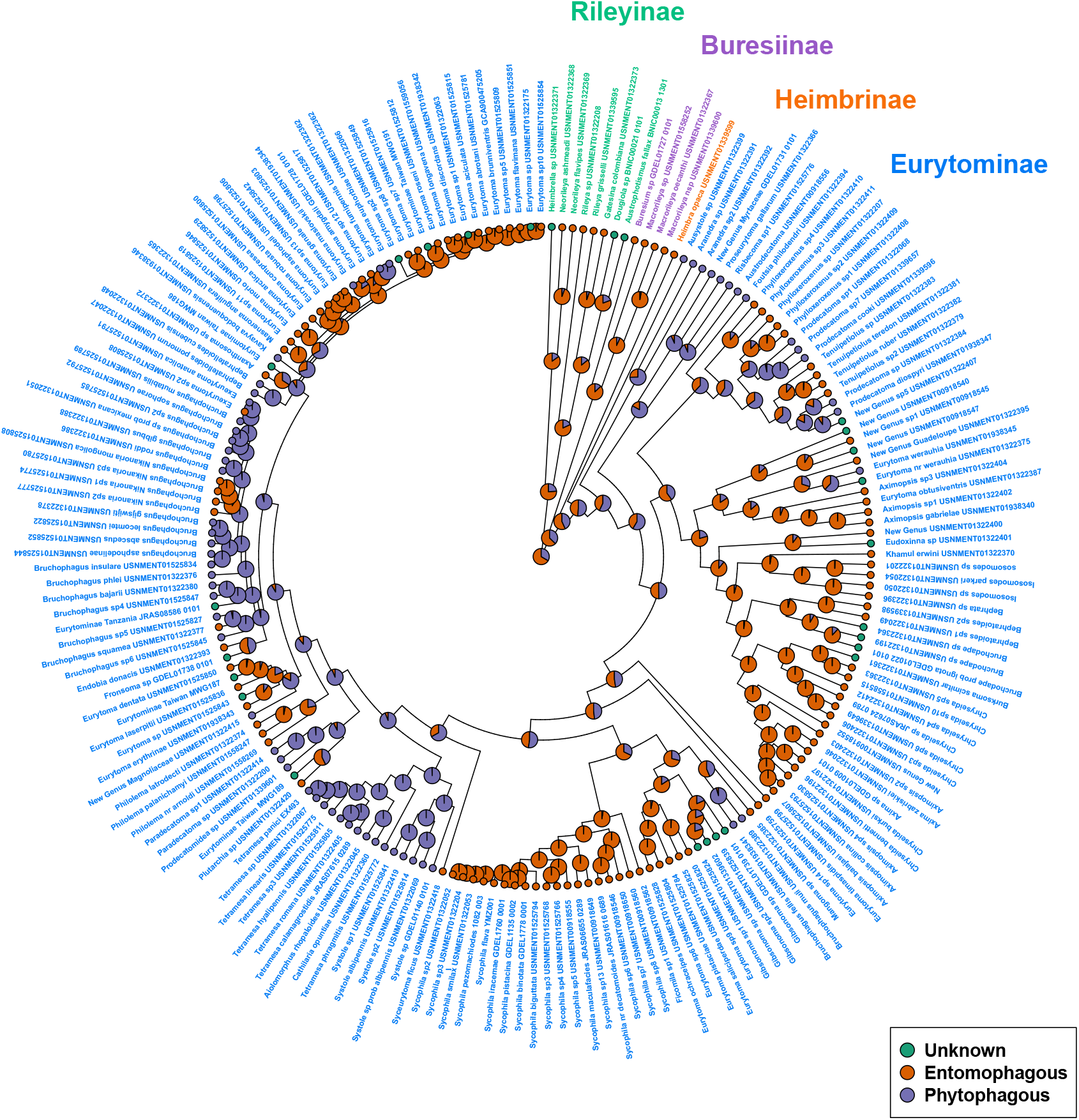
Ancestral state reconstruction of Eurytomidae biology based on the 70p_SWSC matrix, inferred using the ARD model. Green = Unknown Biology, Orange = Parasitoid, Purple = Phytophagous.

When mapping the phytophagous clades on the tree, the propensity for secondary phytophagy appears a number of times according to our ancestral state reconstruction analysis. Phytophagy (including seed feeders and gall formers) occurs in the earliest branching clades of eurytomines in our study, which are mostly Neotropical. However, it is difficult to assess the true biology of many of the eurytomine species as their larvae can be parasitoids that attack their hosts in concealed habitats (galls, twigs or fruit) making it difficult to establish the trophic relationships. In addition, it is known that larvae of some species start their development as parasitoids, and once the host is consumed continue their growth by feeding on plant tissues (La Salle, 2005). Finally in some species, such as *Sycophila* developing at the expense of an epichrysomallid in sycones of figs, are known to be inquilines. Thus, masked phytophagy may be widespread and detailed studies are needed to determine the feeding habits for many groups of eurytomine wasps.

The adaptation from an entomophagous ancestor to phytophagy indeed presupposes that the larvae are able to develop with a completely different diet and have the relevant esterases for this purpose. The switch to full herbivory could be facilitated by the intermediate entomophytophagy lifestyle, as the larvae were already spending much of their life embedded in plant tissues (La Salle, 2005; Tooker & Giron, 2020). Eurytomine species are capable of galling different parts of the plants (roots, twigs, flowers or maturing fruit); it is possible that the larva exudes chemicals mimicking plant hormones for inducing a rapid cell division. Gall induction has evolved multiple times within Chalcidoidea, including within Eurytomidae itself (La Salle, 2005; Tooker & Giron, 2020; Cruaud et al., 2024). To date no comparative studies have been done on gall induction across Chalcidoidea and/or Hymenoptera, so it is unknown how similar the gall induction process by Eurytomidae is compared to the relatively better studied gall wasps and allies (superfamily Cynipoidea). The galls of chalcidoids, including eurytomids, are relatively simple and do not have many defensive traits compared to galls induced by cynipoids, yet it occurs across a much wider variety of host plants ranging from monocot to dicot (La Salle, 2005). Similarly, many eurytomid species are capable of feeding on a variety of seeds, some of which have toxic chemicals (e.g., Rosaceae, Fabaceae), but little is known whether they can sequester these toxic chemicals for their protection against seed-eaters, and whether the detoxification of different chemistry of the host plant has led to the diversification of eurytomids.

Blaimer et al. (2023) examined the link between innovations and diversification in Hymenoptera, including secondary phytophagy. The authors recover a positive diversification rate in the Cynipidae and the Eurytominae, both lineages that have repeated evolution of secondary phytophagy. It is thus possible that the plasticity of Eurytominae larvae for the diet provides an advantage by allowing new niches to be explored, increasing their radiation rate and diversity. A large number of undescribed eurytomines have been recorded from various plants in tropical forest habitats (GD, unpublished data). Based on these observations, we can anticipate that the actual number of species is much higher than currently recognized. The predominantly endophytic lifestyle has no doubt led to a very successful global distribution and rapid diversification of eurytomine wasps, as shown by the short branch lengths in our analyses. Future studies with more detailed taxon sampling of eurytomines could focus on, for example, whether sister group pairs with different life history strategies and/or hosts with dissimilar chemicals have different diversification rates.

## Supporting information

Supplemental Taxonomy

## Acknowledgements

We thank Hossein Lotfalizadeh (Iranian Research Institute of Plant Protection), Chi-Feng Lee (Taiwan Agricultural Research Institute), and Pieter Scott/Jessa Thurman (University of Queensland) for providing additional specimens used in this work. Dietrich Gotzek, Jeffrey Sosa-Calvo, Michael W. Lloyd, Jignasha Rana, Tamara Spasojevic, R. Luke Kresslein, and the LAB facilities of the National Museum of Natural History for providing time and expertise for the UCE library preparation. We acknowledge the Smithsonian Institution High Performance Cluster (https://doi.org/10.25572/SIHPC) for providing computational resources and support that have contributed to the research results reported in this publication. YMZ was funded partially by the Oak Ridge Institute for Science and Education (ORISE) fellowship during this work. The sequencing effort was supported in part by a Smithsonian Institute for Biodiversity Genomics and Global Genome Initiative grant. Mention of trade names or commercial products in this publication is solely for the purpose of providing specific information and does not imply recommendation or endorsement by the USDA. USDA is an equal opportunity provider and employer.

## Data Availability

Clean FASTQ files for each sample newly generated in this study are uploaded onto the SRA database under Bioproject PRJNA1217593, *COI* barcodes are uploaded onto GenBank under accession numbers PV031733–PV031868. All UCE matrices and codes used for analyses are uploaded onto Dryad https://doi.org/10.5061/dryad.3bk3j9kx3.

## Notes

### Competing Interest Statement

The authors have declared no competing interest.

### Summary of Updates

Reanalyzed the data with more stringent filtering, fixed typos and figures.

## References

Beaulieu, J.M., O’Meara, B.C., & Donoghue, M.J. (2013) Identifying hidden rate changes in the evolution of a binary morphological character: the evolution of plant habit in campanulid angiosperms. Systematic Biology, 62(5), 725–737. Available from: 10.1093/sysbio/syt034

Benton, M.J., Wilf, P. & Sauquet, H. (2022) The angiosperm terrestrial revolution and the origins of modern biodiversity. New Phytologist, 233(5), 2017–2035. Available from: 10.1111/nph.17822

Blaimer, B.B., Santos, B.F., Cruaud, A., Gates, M.W., Kula, R.R., Mikó, I. et al. (2023) Key innovations and the diversification of Hymenoptera. Nature communications, 14(1), 1212. Available from: 10.1038/s41467-023-36868-4

Bolger, A.M., Lohse, M., & Usadel, B. (2014) Trimmomatic: a flexible trimmer for Illumina sequence data. Bioinformatics, 30(15), 2114–2120. Available from: 10.1093/bioinformatics/btu170

Borowiec, M.L. (2016) AMAS: a fast tool for alignment manipulation and computing of summary statistics. PeerJ, 4, e1660. Available from: 10.7717/peerj.1660

Borowiec, M.L. (2019a) Spruceup: fast and flexible identification, visualization, and removal of outliers from large multiple sequence alignments. Journal of Open Source Software, 4(42), 1635. Available from: 10.21105/joss.01635

Borowiec, M.L. (2019b) Convergent evolution of the army ant syndrome and congruence in big-data phylogenetics. Systematic Biology, 68(4), 642–656. Available from: 10.1093/sysbio/syy088

Branstetter, M.G., Danforth, B.N., Pitts, J.P., Faircloth, B.C., Ward, P.S., Buffington, M.L. et al. (2017) Phylogenomic insights into the evolution of stinging wasps and the origins of ants and bees. Current Biology, 27(7), 1019–1025. Available from: 10.1016/j.cub.2017.03.027

Branstetter, M.G., Longino, J.T., Ward, P.S., Faircloth, B.C., & Price, S. (2017) Enriching the ant tree of life: enhanced UCE bait set for genome-scale phylogenetics of ants and other Hymenoptera. Methods in Ecology and Evolution, 8(6), 768–776. Available from: 10.1111/2041-210x.12742

Brasero, N. and Martin, N. (2009) Systématique des Chalcidoidea de l’ambre de l’Oise. Mémoire en Sciences Biologiques, Université de Mons-Hainaut, 1–96 pp.

Campbell, B., Heraty, J., Rasplus, J.-Y., Chan, K., Steffen-Campbell, J., & Babcock, C. (2000) Molecular systematics of the Chalcidoidea using 28S-D2 rDNA. In A. Austin & M. Dowton (Eds.), Hymenoptera Evolution, Biodiversity and Biological Control. pp. 59–73. CSIRO Publishing.

Campos-Moreno, D.F., Gates, M.W., Zhang, Y.M., Pérez-Lachaud, G., Dyer, L.A., Whitfield, J.B. et al. (2022) Aximopsis gabrielae sp. nov.: a gregarious parasitoid (Hymenoptera: Eurytomidae) of the skipper Quadrus cerialis (Lepidoptera: Hesperiidae) feeding on Piper amalago in southern Mexico. Journal of Natural History, 56(1-4), 173–189. Available from: 10.1080/00222933.2022.2025940

Castresana, J. (2000) Selection of conserved blocks from multiple alignments for their use in phylogenetic analysis. Molecular Biology and Evolution, 17(4), 540–552. Available from: 10.1093/oxfordjournals.molbev.a026334

Chen, Y., Xiao, H., Fu, J., & Huang, D.-W. (2004) A molecular phylogeny of eurytomid wasps inferred from DNA sequence data of 28S, 18S, 16S, and COI genes. Molecular Phylogenetics and Evolution, 31(1), 300–307. Available from: 10.1016/s1055-7903(03)00282-3

Cruaud, A., Delvare, G., Nidelet, S., Sauné, L., Ratnasingham, S., Chartois, M. et al. (2021) UltralJConserved Elements and morphology reciprocally illuminate conflicting phylogenetic hypotheses in Chalcididae (Hymenoptera, Chalcidoidea). Cladistics, 37, 1–35. Available from: 10.1111/cla.12416

Cruaud, A., Nidelet, S., Arnal, P., Weber, A., Fusu, L., Gumovsky, A. et al. (2019) Optimized DNA extraction and library preparation for minute arthropods: Application to target enrichment in chalcid wasps used for biocontrol. Molecular Ecology Resources, 19(3): 702–710. Available from: 10.1111/1755-0998.13006

Cruaud, A., Rasplus, J.-Y., Zhang, J., Burks, R., Delvare, G., Fusu, L. et al. (2024) The Chalcidoidea bush of life: evolutionary history of a massive radiation of minute wasps. Cladistics, 40, 34–63. Available from: 10.1111/cla.12561

Faircloth, B.C. (2013) Illumiprocessor: a trimmomatic wrapper for parallel adapter and quality trimming. Available from: 10.6079/J9ILL

Faircloth, B.C. (2015) PHYLUCE is a software package for the analysis of conserved genomic loci. Bioinformatics, 32(5), 786–788. Available from: 10.1093/bioinformatics/btv646

Faircloth, B.C., Branstetter, M.G., White, N.D., & Brady, S.G. (2015) Target enrichment of ultraconserved elements from arthropods provides a genomic perspective on relationships among Hymenoptera. Molecular Ecology Resources, 15(3), 489–501. Available from: 10.1111/1755-0998.12328

Faircloth, B.C., McCormack, J.E., Crawford, N.G., Harvey, M.G., Brumfield, R.T., & Glenn, T.C. (2012) Ultraconserved elements anchor thousands of genetic markers spanning multiple evolutionary timescales. Systematic Biology, 61(5), 717–726. Available from: 10.1093/sysbio/sys004

Gates, M.W. (2008) species revision and generic systematics of world Rileyinae (Hymenoptera: Eurytomidae) (Vol. 127). Univ of California Press.

Gates, M.W. & Cascante-Marin, A. (2004) A new phytophagous species of Eurytoma (Hymenoptera: Eurytomidae) attacking Werauhia gladioliflora (Bromeliales: Bromeliaceae). Zootaxa, 512(1), 1–10. Available from: 10.11646/zootaxa.512.1.1

Gates, M. & Delvare, G. (2008) A new species of Eurytoma (Hymenoptera: Eurytomidae) attacking Quadrastichus spp. (Hymenoptera: Eulophidae) galling Erythrina spp. (Fabaceae), with a summary of African Eurytoma biology and species checklist. Zootaxa, 1751(1), 1–24. Available from: 10.11646/zootaxa.1751.1.1

Gates, M.W., Delvare, G., & Zhang, Y.M. (2025) Eurytomidae. In J. M. Heraty, Woolley, J. B. (Ed.), Chalcidoidea of the World. CABI Press, Wallingford, UK.

Glenn, T.C., Pierson, T.W., Bayona-Vásquez, N.J., Kieran, T.J., Hoffberg, S.L., Thomas Iv, J.C. et al. (2019) Adapterama II: universal amplicon sequencing on Illumina platforms (TaggiMatrix). PeerJ, 7. Available from: 10.7717/peerj.7786

Guindon, S., Dufayard, J.-F., Lefort, V., Anisimova, M., Hordijk, W., & Gascuel, O. (2010) New algorithms and methods to estimate maximum-likelihood phylogenies: assessing the performance of PhyML 3.0. Systematic Biology, 59(3), 307–321. Available from: 10.1093/sysbio/syq010

Heraty, J.M., Burks, R.A., Cruaud, A., Gibson, G.A., Liljeblad, J., Munro, J. et al. (2013) A phylogenetic analysis of the megadiverse Chalcidoidea (Hymenoptera). Cladistics, 29(5), 466–542. Available from: 10.1111/cla.12006

Hoang, D.T., Chernomor, O., von Haeseler, A., Minh, B.Q., & Le, S.V. (2017) UFBoot2: Improving the ultrafast bootstrap approximation. Molecular Biology and Evolution, msx281. Available from: 10.1093/molbev/msx281

Jafari, S., Müller, B., Rulik, B., Rduch, V., & Peters, R.S. (2023) Another crack in the Dark Taxa wall: a custom DNA barcoding protocol for the species-rich and common Eurytomidae (Hymenoptera, Chalcidoidea). Biodiversity Data Journal, 11, e101998. Available from: 10.3897/BDJ.11.e101998.

Junier, T., & Zdobnov, E.M. (2010) The Newick utilities: high-throughput phylogenetic tree processing in the UNIX shell. Bioinformatics, 26(13), 1669–1670. Available from: 10.1093/bioinformatics/btq243

Kalyaanamoorthy, S., Minh, B.Q., Wong, T.K., von Haeseler, A., & Jermiin, L.S. (2017) ModelFinder: fast model selection for accurate phylogenetic estimates. Nature Methods, 14(6), 587. Available from: 10.1038/nmeth.4285

Katoh, K., Misawa, K., Kuma, K.-i., & Miyata, T. (2002) MAFFT: a novel method for rapid multiple sequence alignment based on fast Fourier transform. Nucleic Acids Research, 30(14), 3059–3066. Available from: 10.1093/nar/gkf436

Katoh, K., & Toh, H. (2008) Recent developments in the MAFFT multiple sequence alignment program. Briefings in Bioinformatics, 9(4), 286–298. Available from: 10.1093/bib/bbn013

Lanfear, R., Calcott, B., Kainer, D., Mayer, C., & Stamatakis, A. (2014) Selecting optimal partitioning schemes for phylogenomic datasets. BMC Evolutionary Biology, 14(1), 1–14. Available from: 10.1186/1471-2148-14-82

Lanfear, R., Frandsen, P.B., Wright, A. M., Senfeld, T., & Calcott, B. (2016) PartitionFinder 2: new methods for selecting partitioned models of evolution for molecular and morphological phylogenetic analyses. Molecular Biology and Evolution, 34(3), 772–773. Available from: 10.1093/molbev/msw260

La Salle, J. (2005) Biology of gall inducers and evolution of gall induction in Chalcidoidea (Hymenoptera: Eulophidae, Eurytomidae, Pteromalidae, Tanaostigmatidae, Torymidae). In A. Raman, C.W. Schaefer, T.M. Withers (Eds.), Biology, Ecology, and Evolution of Gall-inducing Arthropods. pp. 509–537. Science Publishers, Inc., Enfield, NH.

Li, Y., Zhou, X., Feng, G., Hu, H., Niu, L., Hebert, P.D. et al. (2010) COI and ITS2 sequences delimit species, reveal cryptic taxa and host specificity of fig-associated Sycophila (Hymenoptera, Eurytomidae). Molecular Ecology Resources, 10(1), 31–40. Available from: 10.1111/j.1755-0998.2009.02671.x

Lotfalizadeh, H., Delvare, G., Cruaud, A., & Rasplus, J.-Y. (2024) Morphological phylogeny and revision of Sycophila and Ficomila (Hymenoptera: Chalcidoidea, Eurytomidae) associated with Afrotropical fig trees (Moraceae, Ficus). Zootaxa, 5401(1), 1–190. Available from: 10.11646/zootaxa.5401.1.1

Lotfalizadeh, H., Delvare, G., & Rasplus, J.-Y. (2007) Phylogenetic analysis of Eurytominae (Chalcidoidea: Eurytomidae) based on morphological characters. Zoological Journal of the Linnean Society, 151(3), 441–510. Available from: 10.1111/j.1096-3642.2007.00308.x

Matzke, N.J. (2014) Model selection in historical biogeography reveals that founder-event speciation is a crucial process in Island clades. Systematic Biology, 63, 951–970. Available from: 10.1093/sysbio/syu056

Matzke, N.J. (2018) BioGeoBEARS: BioGeography with Bayesian (and likelihood) Evolutionary Analysis with R Scripts. version 1.1.1, published on GitHub on November 6, 2018. Available from: 10.5281/zenodo.1478250

Matzke, N.J. (2022) Statistical comparison of DEC and DEC + J is identical to comparison of two ClaSSE submodels, and is therefore valid. Journal of Biogeography, 49, 1805–1824. Available from: 10.1111/jbi.14346

Minh, B.Q., Schmidt, H. A., Chernomor, O., Schrempf, D., Woodhams, M.D., Von Haeseler, A. et al. (2020a). IQ-TREE 2: New models and efficient methods for phylogenetic inference in the genomic era. Molecular Biology and Evolution, 37(5), 1530–1534. Available from: 10.1093/molbev/msaa131

Minh, B.Q., Hahn, M.W., & Lanfear, R. (2020b) New methods to calculate concordance factors for phylogenomic datasets. Molecular Biology and Evolution, 37(9), 2727–2733. Available from: 10.1093/molbev/msaa106

Munro, J.B., Heraty, J.M., Burks, R.A., Hawks, D., Mottern, J., Cruaud, A. et al. (2011) A molecular phylogeny of the Chalcidoidea (Hymenoptera). PLoS One, 6(11), e27023. Available from: 10.1371/journal.pone.0027023

Naser-Khdour, S., Minh, B.Q., Zhang, W., Stone, E.A., & Lanfear, R. (2019) The prevalence and impact of model violations in phylogenetic analysis. Genome Biology and Evolution, 11(12), 3341–3352. Available from: 10.1093/gbe/evz193

Peris, D., & Condamine, F.L. (2024) The angiosperm radiation played a dual role in the diversification of insects and insect pollinators. Nature Communications, 15(1), 552. Available from: 10.1038/s41467-024-44784-4

Peters, R.S., Niehuis, O., Gunkel, S., Blaser, M., Mayer, C., Podsiadlowski, L. et al. (2018). Transcriptome sequence-based phylogeny of chalcidoid wasps (Hymenoptera: Chalcidoidea) reveals a history of rapid radiations, convergence, and evolutionary success. Molecular Phylogenetics and Evolution, 120, 286–296. Available from: 10.1016/j.ympev.2017.12.005

Prjibelski, A., Antipov, D., Meleshko, D., Lapidus, A., & Korobeynikov, A. (2020) Using SPAdes de novo assembler. Current Protocols in Bioinformatics, 70(1), e102. Available from: 10.1002/cpbi.102

Puttick, M.N. (2019) MCMCtreeR: functions to prepare MCMCtree analyses and visualize posterior ages on trees. Bioinformatics, 35(24), 5321–5322. Available from: 10.1093/bioinformatics/btz554

R Core Team. (2024) R: A language and environment for statistical computing. Available from: https://www.R-project.org/

Rambaut, A., Drummond, A.J., Xie, D., Baele, G., & Suchard, M.A. (2018) Posterior summarization in Bayesian phylogenetics using Tracer 1.7. Systematic Biology, 67(5), 901. Available from: 10.1093/sysbio/syy032say

Rasplus, J.-Y., Heraty, J.M., Burks, R.A., Krogman, L., Ulmer, J.M. & Cruaud, A. (2025) Fossils of Chalcidoidea. In J. M. Heraty, Woolley, J. B. (Ed.), Chalcidoidea of the World. CABI Press, Wallingford, UK.

Ree, R.H., & Smith, S.A. (2008) Maximum-likelihood inference of geographic range evolution by dispersal, local extinction, and cladogenesis. Systematic Biology, 57, 4–14. Available from: 10.1080/10635150701883881

Revell, L.J. (2024) phytools 2.0: an updated R ecosystem for phylogenetic comparative methods (and other things). PeerJ, 12, e16505. Available from: 10.7717/peerj.16505

Ronquist, F. (1997) Dispersal-vicariance analysis: a new approach to the quantification of historical biogeography. Systematic Biology, 46, 195–203. Available from: 10.1093/sysbio/46.1.195

Sayyari, E., & Mirarab, S. (2016) Fast coalescent-based computation of local branch support from quartet frequencies. Molecular Biology and Evolution, 33(7), 1654–1668. Available from: 10.1093/molbev/msw079

Stage, G.I., & Snelling, R.R. (1986) The subfamilies of Eurytomidae and systematics of the subfamily Heimbrinae (Hymenoptera: Chalcidoidea). Contributions in Science. Natural History Museum of Los Angeles County (375).

Stamatakis, A. (2006) RAxML-VI-HPC: maximum likelihood-based phylogenetic analyses with thousands of taxa and mixed models. Bioinformatics, 22(21), 2688–2690. Available from: 10.1093/bioinformatics/btl446

Tagliacollo, V.A., & Lanfear, R. (2018) Estimating improved partitioning schemes for ultraconserved elements. Molecular Biology and Evolution, 35(7), 1798–1811. Available from: 10.1093/molbev/msy069

Tooker, J.F. & Giron, D. (2020) The evolution of endophagy in herbivorous insects. Frontiers in Plant Science, 11, 581816. Available from: 10.3389/fpls.2020.581816

UCD Community. (2025) Universal Chalcidoidea Database https://ucd.chalcid.org x[accessed 1/1/2025)

Wang, H.-C., Minh, B.Q., Susko, E., & Roger, A.J. (2018) Modeling site heterogeneity with posterior mean site frequency profiles accelerates accurate phylogenomic estimation. Systematic Biology, 67(2), 216–235. Available from: 10.1093/sysbio/syx068

Wang, L.G., Lam, T.T.Y., Xu, S., Dai, Z., Zhou, L., Feng, T., et al. (2020). Treeio: an R package for phylogenetic tree input and output with richly annotated and associated data. Molecular Biology and Evolution, 37(2), 599–603. Available from: 10.1093/molbev/msz240

Yang, Z. (2007) PAML 4: phylogenetic analysis by maximum likelihood. Molecular Biology and Evolution, 24(8), 1586–1591. Available from: 10.1093/molbev/msm088

Yu, G., Smith, D.K., Zhu, H., Guan, Y., & Lam, T.T.Y. (2017) ggtree: an R package for visualization and annotation of phylogenetic trees with their covariates and other associated data. Methods in Ecology and Evolution, 8(1), 28–36. Available from: 10.1111/2041-210X.12628

Zerova, M.D. (1988) The main trends of the evolution and system of chalcids of the family Eurytomidae (Hymenoptera, Chalcidoidea). Entomologicheskoe Obozrenie, 67, 649–674.

Zhang, C., Rabiee, M., Sayyari, E., & Mirarab, S. (2018) ASTRAL-III: polynomial time species tree reconstruction from partially resolved gene trees. BMC Bioinformatics, 19(6), 153. Available from: 10.1186/s12859-018-2129-y

Zhang, J., Lindsey, A.R. I., Peters, R.S., Heraty, J.M., Hopper, K.R., Werren, J.H. et al. (2020). Conflicting signal in transcriptomic markers leads to a poorly resolved backbone phylogeny of chalcidoid wasps. Systematic Entomology, 45, 783–802. Available from: 10.1111/syen.12427

Zhang, Y.M., Bossert, S. & Spasojevic, T. (2025) Evolving perspectives in Hymenoptera systematics: Bridging fossils and genomes across time. Systematic Entomology, 50(1), 1–31. Available from: 10.1111/syen.12645

Zhang, Y.M., Gates, M.W. & Shorthouse, J.D. (2014) Testing species limits of Eurytomidae (Hymenoptera) associated with galls induced by Diplolepis (Hymenoptera: Cynipidae) in Canada using an integrative approach. The Canadian Entomologist, 146(3), 321–334. Available from: 10.4039/tce.2013.70

Zhang, Y.M., Gates, M.W., Silvestre, R. & Scarpa, M. (2021) Description of Kavayva, gen. nov., (Chalcidoidea, Eurytomidae) and two new species associated with Guarea (Meliaceae), and a review of New World eurytomids associated with seeds. Journal of Hymenoptera Research, 86, 101–121. Available from: 10.3897/jhr.86.71309

Zhang, Y.M., Sheikh, S.I., Ward, A.K.G., Forbes, A.A., Prior, K.M., Stone, G.N. et al. (2022). Delimiting the cryptic diversity and host preferences of Sycophila parasitoid wasps associated with oak galls using phylogenomic data. Molecular Ecology, 31(16), 4417–4433. Available from: 10.1111/mec.16582

Zhang, Y.M., Williams, J.L., & Lucky, A. (2019) Understanding UCEs: A comprehensive primer on using Ultraconserved Elements for arthropod phylogenomics. Insect Systematics and Diversity, 3(5), 3. Available from: 10.1093/isd/ixz016

