## Supplementary figures and images for "Phasing in and out of phytophagy: phylogeny and evolution of the family Eurytomidae (Hymenoptera: Chalcidoidea) based on Ultraconserved Elements"

### Supplemental Figure 1

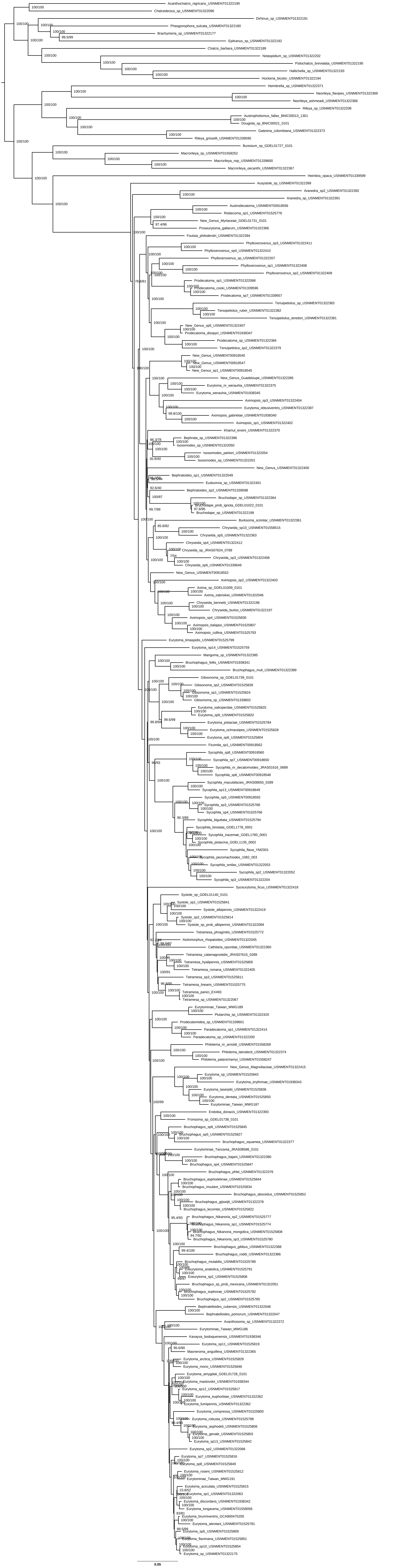

### Supplemental Figure 2

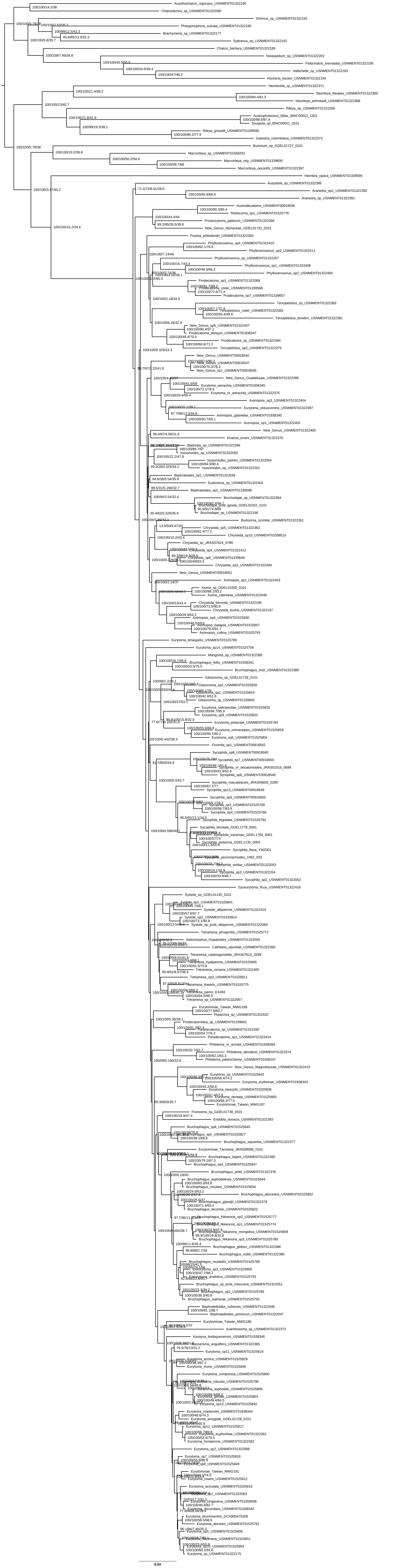

### Supplemental Figure 3

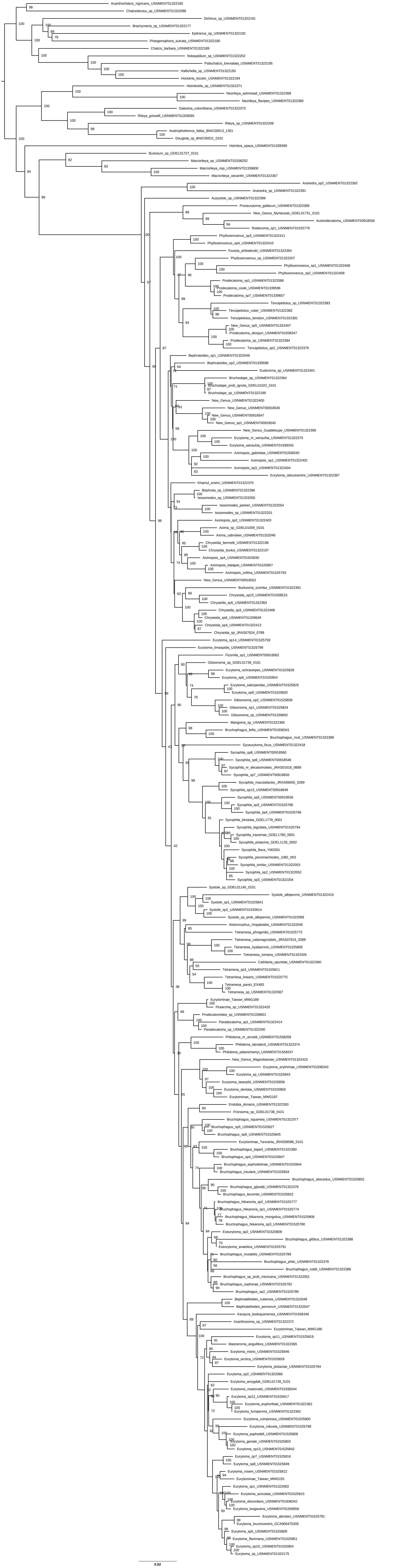

### Supplemental Figure 4

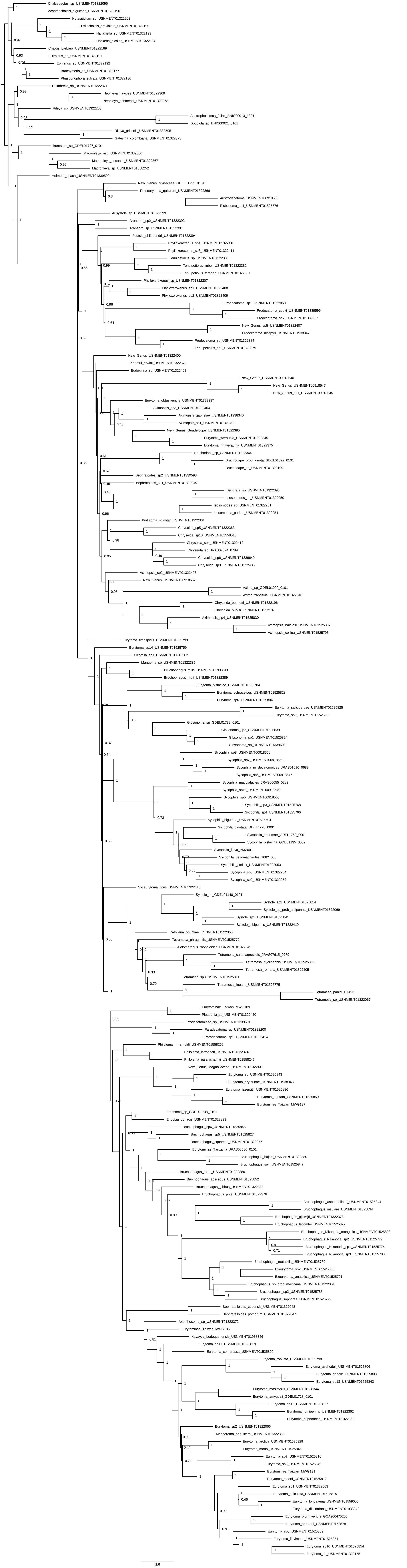

### Supplemental Figure 5

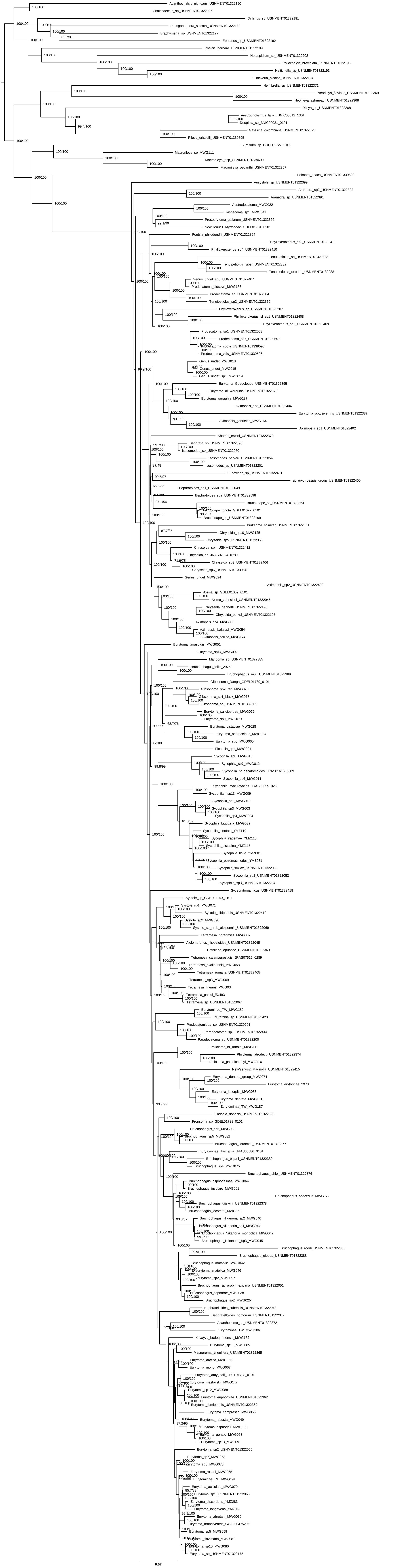

### Supplemental Figure 6

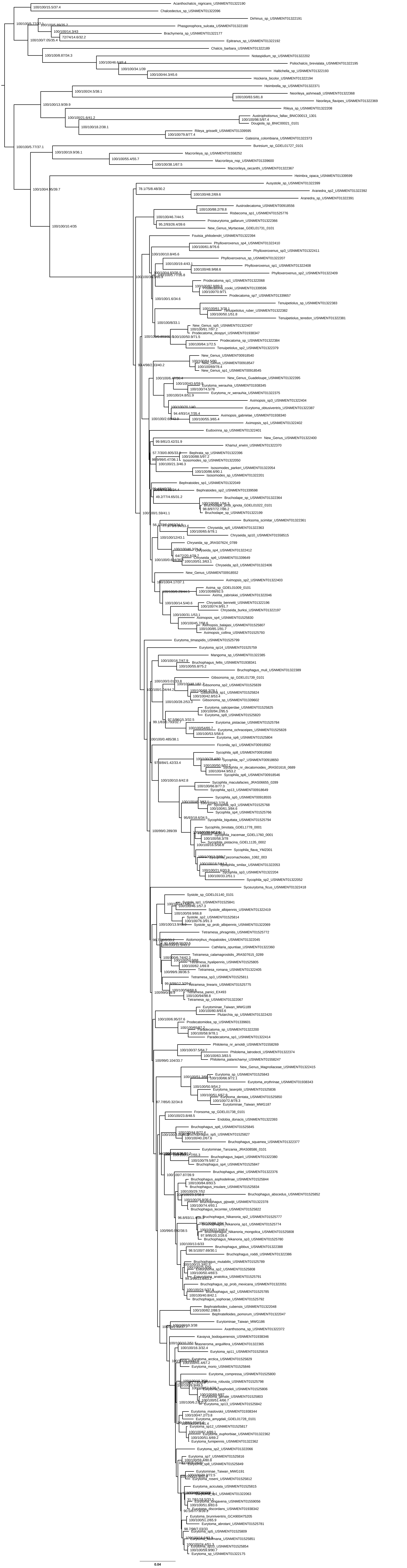

### Supplemental Figure 7

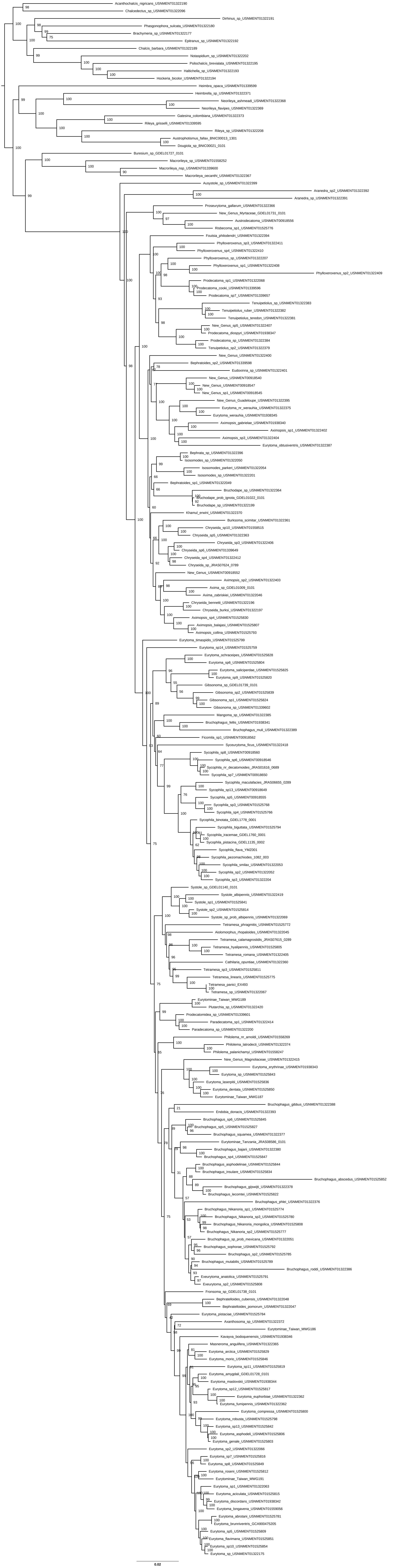

### Supplemental Figure 8

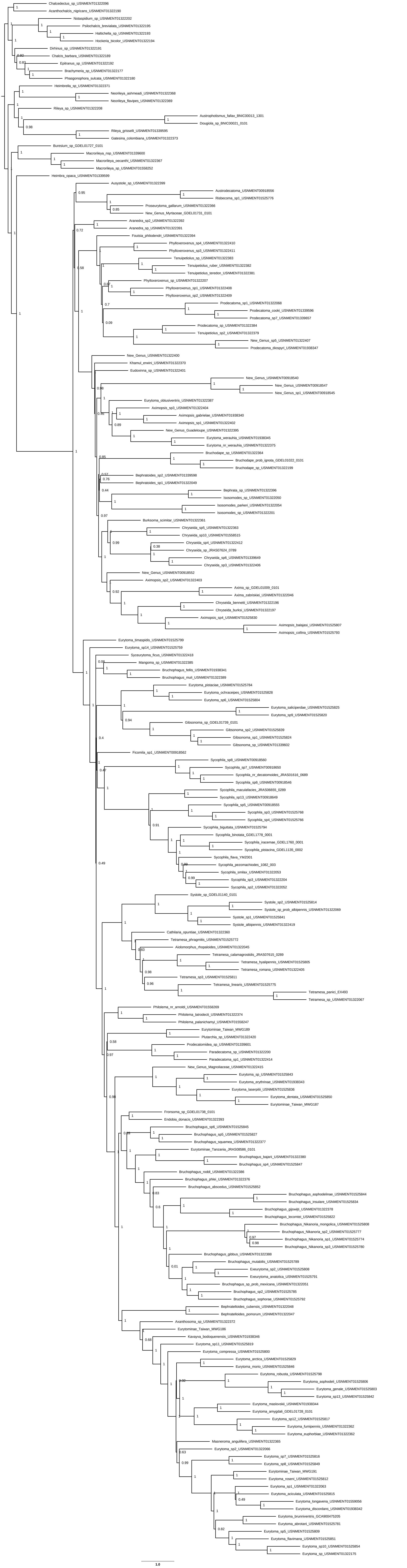

### Supplemental Figure 9

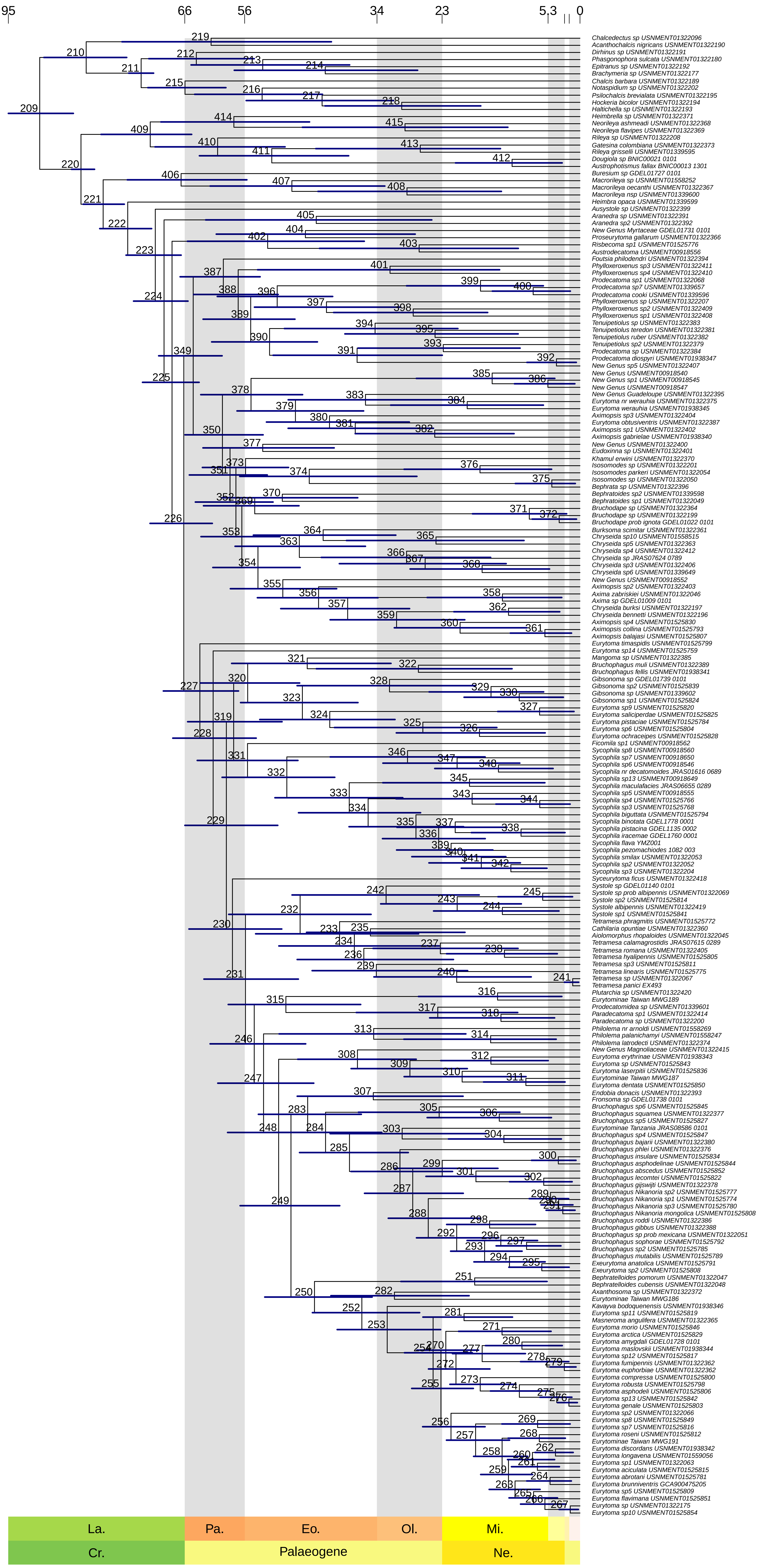

### Supplemental Figure 10

BioGeoBEARS DIVALIKE on Eurytomidae M1_constrained
ancstates: global optim, 5 areas max. d=0.0031; e=0; j=0; LnL=−282.30

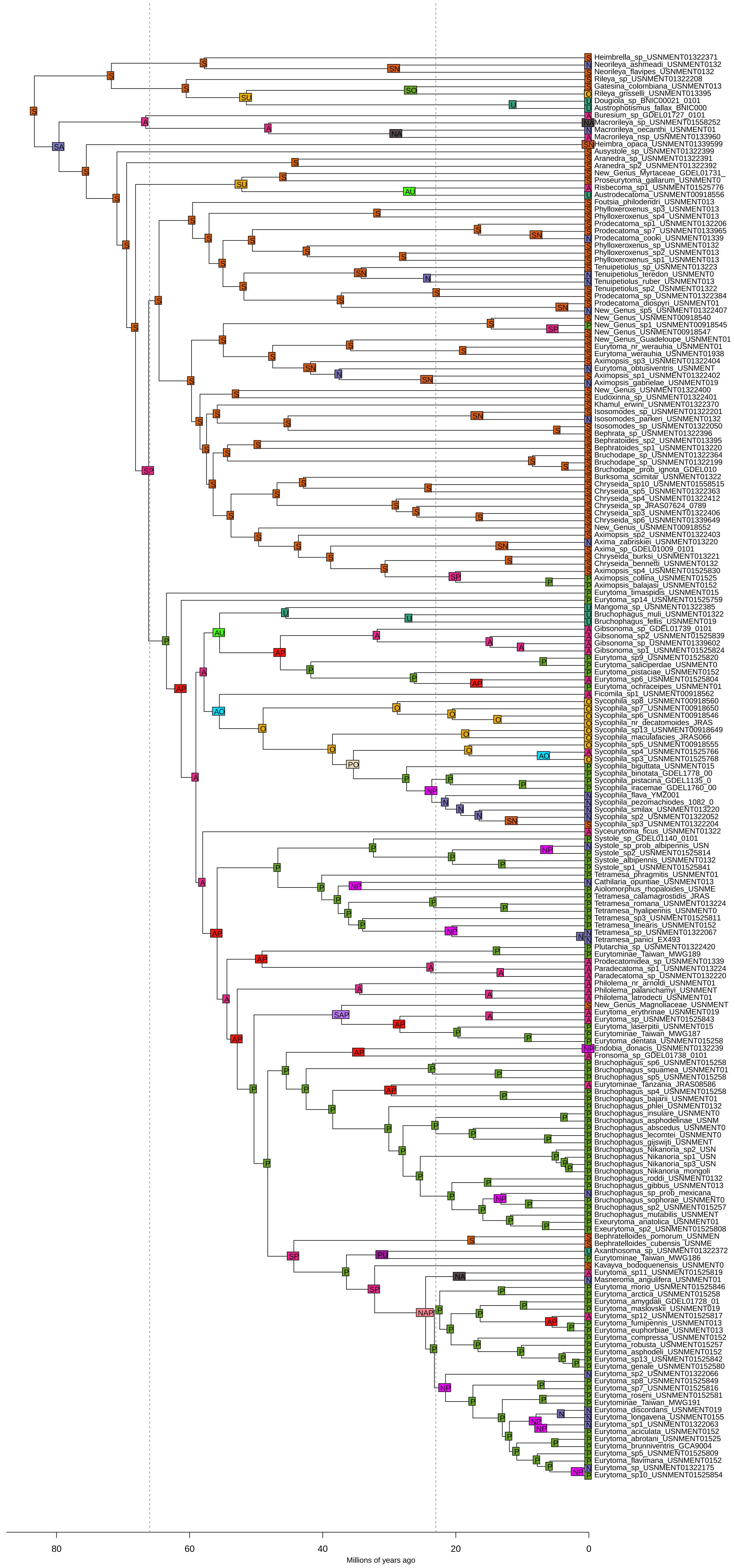
