## Supplemental Taxonomy for "Phasing in and out of phytophagy: phylogeny and evolution of the family Eurytomidae (Hymenoptera: Chalcidoidea) based on Ultraconserved Elements"

**Missing taxa in the sampling**

Subfamily Rileyinae (1 sp.)

*Platyrileya* Burks, 1971; type species *Platyrileya cururipe* Burks, 1971(1 sp.)

Subfamily Heimbrinae (2 spp.)

*Symbra* Stage & Snelling, 1986; type species *Symbra cordobensis* Stage & Snelling, 1986 (2 sp)

Subfamily Eurytominae (21 genera missing but 46 spp. only)

*Axanthosomella* Narendran, 2001; type species *Axanthosomella gadagkari* Narendran, 2001 (1 sp.) [probably a junior synonym of *Tetramesa* according to original description]

*Banyoma* Burks, 1971; type species *Banyoma philippinensis* Burks (2 spp.)

*Camponotophilus* Gates, 2012; type species *Camponotophilus* delvarei Gates, 2012 (1 sp.)

*Eurytomocharis* Ashmead, 1888; type species *Eurytomocharis minuta* Ashmead, 1888 (18 spp.)

*Giraultoma* Boućek, 1988; type species *Xanthosoma pulchricorpus* (Girault, 1915) (2 spp.)

*Hexeurytoma* Dodd, 1917; type species *Hexeurytoma grandis* Dodd, 1917 (3 spp.)

*Homodecatoma* Liao, 1979; type species *Homodecatoma mallotae* Liao, 1979 (1 sp.)

*Houstonia zani* Boućek, 1988; type species *Houstonia zani* Boućek,1988 (1 sp.)

*Neobephrata* Narendran & Padmasenan, 1989; type species *Neobephrata petiolata* Narendran & Padmasenan, 1989 (1 sp.)

*Neoeurytomaria* Narendran, 1994; type species *Neoeurytomaria subbaraoi* Narendran, 1994 (1 sp.)

*Philippinoma* Narendran, 1994; type species *Philippinoma auratofronta* Narendran, 1994 (2 spp.)

*Phleudecatoma* Yang, 1996; type species *Phleudecatoma platycladi* Yang, 1996 (2 spp.)

*Pseudotetramesa* Kalina,1970; type species *Pseudotetramesa doksensis* Kalina, 1970 (1 sp.)

*Ramanuja* Narendran, 1989; type species *Ramanuja swarnamus* Narendran, 1989 (1 sp.)

*Ramdasoma* Narendran, 1994; type species *Ramdasoma peethodaris* Narendran,1994 (3 spp.)

*Stigmeurytoma* Bouček, 1988; type species *Eurytoma eucalypti* Ashmead, 1900 (1 sp.)

*Subbaella* Narendran, 1994; type species *Subbaella negriensis* Narendran,1994 (1 sp.)

*Systolema* Narendran, 1994; type species *Systolema hayati* Narendran, 1994 (1 sp.)

*Tetramesella* Zerova, 1974; *Tetramesella luppovae* Zerova, 1974 (1 sp.)

*Townosema* Narendran, 1994; type species *Townosema taiwanicus* Narendran, 1994 (1 sp.)

*Zerovella* Narendran & Sheela, 1994; type species *Zerovella taiwanica* Narendran & Sheela, 1994 (1 sp.)

**Detailed taxonomic discussion of subfamily Eurytominae**

The results are largely excellent, as we successfully retrieve many genera—both already described and others that are undescribed or currently considered species groups—previously identified by Lotfalizadeh *et al.* (2007). Thus, the molecular results largely align with those obtained through morphological analyses. However, the same issue persists: the overall phylogenetic pattern remains unresolved, with the backbone of the tree unsolved except for the basal clades (*Ausystole*, *Aranedra*, *Proseurytoma*, *Risbecoma*, and *Austrodecatoma*). Interestingly, these basal clades exclusively consist of phytophagous species, including gall formers.

**Clade *Phylloxeroxenus***

Within the clade *Phylloxeroxenus*, the genera *Phylloxeroxenus* and *Prodecatoma* appear to be polyphyletic, despite the presence of several derived characters supporting their monophyly as suggested by Lotfalizadeh *et al.* (2007). However, *Phylloxeroxenus* exhibits significant morphological variation, indicating that multiple distinct taxa may be involved (GD, pers. observ.). Additionally, the backbone of the clade remains unresolved, with the basal nodes positioned extremely close to one another. As a result, it is currently impossible to draw firm conclusions regarding the potential paraphyly of *Phylloxeroxenus* and *Prodecatoma*.

**Clade *Chryseida***

Generally speaking, the backbone of the clade *Chryseida* is not resolved, suggesting rapid radiation; thus, the relationships between the different sets included here are somewhat hypothetical. *Eurytoma werauhia* was erroneously found closely related to *Paradecatoma* by

Lotfalizadeh *et al.* (2007); it is actually in the same genus as New_Genus_Guadeloupe_USNMENT01322395 within our current study. *Aximopsis*_sp1_USNMENT01322402 and the recently described *Aximopsis gabrielae* do not belong to *Aximopsis* due to their distinctive features on the head and antenna. Similarly, the closely related *Eurytoma obtusiventris* also differs from *Aximopsis* as it lacks the latter's derived traits, particularly the deep subventral pit of the prepectus.

The genus *Chryseida* is split into three sets, appearing paraphyletic relative to *Burksoma*. Such a relationship is surprising given the distinct morphology of *Chryseida*; however, *Burksoma* is retrieved on a long branch, making this relationship doubtful. The specimen New_Genus_USNMENT00918552, collected from Peru, does not fit any current generic concept and requires the erection of a new genus to accommodate it. *Chryseida bennetti* Burks and *C. burksi* Zerova were included in this genus due to their very slight metallic reflections. Nevertheless, they share all the derived traits of *Aximopsis sensu largo* and are thus congeneric with it. These two species also lack the spherical or tongue-like projection of the ventral rim of the foramen magnum seen in other *Chryseida* species identified by Lotfalizadeh *et al.* (2007) (MWG, pers. observ.).

**Clade *Eurytoma verticillata* species group**

The placement of *Eurytoma*_sp14_USNMENT01525759, far from the genus *Eurytoma sensu stricto*, is quite surprising. In fact the *verticillata* group, together with *Axanthosoma* and *Masneroma*, exhibits the characteristic postgenal lamina of *Eurytoma*, *e.g.* diverging from the plane of the postgena and thus visible from the side. In addition the results are unstable regarding the position of the relevant specimen (*Eurytoma* sp14) on the tree: in a previous tree it merges on a node much closer to *Eurytoma*. Thus additional data are required to accurately determine the position of the *Eurytoma verticillata* group relative to the main clade of *Eurytoma*.

**Clade *Eurytoma aspila* species group**

It is represented in the present study by the single specimen *Eurytoma_timaspidis*_USNMENT01525799, appearing as the sister group of all the terminal clades of Eurytominae starting with the *Sycophila* clade. The group thus emerges far from the node supporting *Eurytoma s.s.*; in fact species of the *aspila* group lack the postgenal lamina retrieved in *Eurytoma.* Their body is bicolored (yellow + black or dark brown) and they show very deep notauli. They all inhabit the Palearctic Region and were reared from cynipid galls on Asteraceae.

**Clade *Sycophila***

The relationship between the two Australasian *Bruchophagus* species associated with *Citrus* (*Bruchophagus fellis* and *B. muli*) aligns with their morphology, biogeography, and biology. It was anticipated that these *Bruchophagus* species do not fit with the rest of the genus but were included due to their phytophagous habit. The erection of a new genus is therefore necessary for the taxonomic placement of this pair, particularly considering the pest status of *B. fellis* in Australia.

The pair of *Eurytoma* species (including *E. saliciperdae*) reared from galls of *Pontania* (Tenthredinidae) on *Salix* evidently do not belong to *Eurytoma*, as they fall well outside the main clade of the genus. They were named as such based on the current official nomenclature. These species were mistakenly included in *Mangoma* by Lotfalizadeh et al. (2007) due to an incorrect phylogenetic placement. *Eurytomocharis* Ashmead, based on the morphology of its type species *Eurytoma minuta* Ashmead (later renamed *E. ashmeadi* (Peck) due to homonymy), appears to be the best fit for them.

*Eurytoma pistaciae* Rondani, *E. ochraceipes* Kalina, and *Eurytoma*_sp6_USNMENT01525804 belong to the *pistaciae* species group, which is easily recognizable by the presence of long pegs on the dorsal side of the metatibia. This group is restricted to the Old World but is much more diverse in tropical regions. The reared specimens emerged from galls or fruit infested by various holometabolous insects (GD, pers. observ.). For the same reasons mentioned above, their taxonomic placement requires the erection of a new genus.

Finally, the diverse genus *Sycophila* Walker is confirmed as monophyletic and appears as the sister group to *Ficomila* Bouček, in agreement with Lotfalizadeh *et al.* (2007) and Lotfalizadeh *et al.* (2024).

### Genus *Syceurytoma*

*Syceurytoma ficus* occupies an isolated branch between two large clades, but the distance between the nodes separating it from the *Systole* clade is relatively short. While its placement remains somewhat uncertain, it is recognized as a distinct monotypic genus.

### Clade *Systole*

Lotfalizadeh *et al.* (2007) reconstructed a similar topology to the present study for this group, with the genus *Systole* as the sister group to the clade comprising *Tetramesa*, *Cathilaria*, and *Aiolomorphus*. However, the authors did not formally synonymize these genera. *Tetramesa* Walker, *Cathilaria* Burks, and *Aiolomorphus* Walker all form galls in twigs or flowers of Poaceae, making such synonymies consistent with their shared biology. This also resolves the paraphyly previously observed in *Tetramesa*. Consequently, the four *Cathilaria* species (*C. certa*, *C. globiventris*, *C. opuntiae*, and *C. rigidae*) and the monotypic *Aiolomorphus rhopaloides* are formally synonymized within *Tetramesa* **syn. nov.**.

### Clade *Plutarchia*

*Plutarchia* was reared from small flies infesting fruit (Bouček, 1988), while *Paradecatoma* was found infesting seeds of various plants (*Cordia* L., *Magnolia* L. spp.) (Yrgu & Delvare, 2019).

Based on its morphology, characterized by a squat body and uncompressed gaster, *Prodecatomidea* is also likely phytophagous (GD, pers. observ.).

**Genus *Philolema***

*Philolema* is retrieved as monophyletic. *Philolema* aff. *arnoldi* is likely a secondary parasitoid of Lepidoptera, while the other species are known to parasitize the eggs of spiders. The present results confirm those of Lotfalizadeh *et al.* (2007).

**Clade *dentata* Group**

The *Eurytoma dentata* species group was used by default, following the present nomenclature. Most species included here (except possibly New_Genus_Magnoliaceae_USNMENT01322415) belong to the same undescribed genus. *E. dentata* Mayr, *E. laserpitii* Mayr, and a few other undescribed species were reared from cecidomyiid flies galling various dicots. The group is present at least in the West Palearctic and Afrotropical regions.

**Clade *Endobia***

The *Endobia* clade includes only the pair *Endobia* Erdös and *Fronsoma* Narendran. The specimen of the latter genus comes from the Afrotropical region, marking a new record for the continent. *Endobia* is known from Southern Europe and India; both species were reared from twigs of gramineous plants, with the first species acting as a parasitoid of bostrichid beetle (*Dinoderus* Stephens) that mines the twigs. The *Fronsoma* specimen in this study was collected in Cameroon while foraging on a dead trunk infested by xylophagous beetles.

**Clade *Bruchophagus***

The *Bruchophagus* clade consists of the majority of *Bruchophagus* Ashmead, including its type species *B. borealis* Ashmead, a species morphologically close to *B. gibbus* Boheman and *B. roddi* Gussakovskyi, both of which are included in the present sampling. These species belong to the large *gibbus* species group, which comprises species that infest seeds of leguminous plants. The topology of the tree follows the partition into species groups as hypothesized by Lotfalizadeh *et al.* (2007), with the following exceptions: 1) The *borealis* group, which was mixed with the species group infesting Asphodelaceae; 2) The *bajarii* species group, which branched outside *Bruchophagus* but merges here on a sub-basal branch. In the present tree, a basal set represents the *squamea* species group. *Bruchophagus phlei*, still classified under *Eurytoma* by Zerova (2010), is the only species representing this group here.

A set of species reared from seeds of Asphodelaceae forms a single branch (*B. lecomtei* Delvare, *B. gijswijti* Askew & Ribes, *B. abscedus* Askew, *B. insulare* Delvare, *B. asphodelinae* Askew & Stojanova). These species are equivalent to the recently erected genus *Parabruchophagus* Zerova (formally a subgenus of *Bruchophagus*), which also develops in the seeds of Asphodelaceae (*Eremurus* spp.) in central Asia (Zerova 2011). Therefore, the genus *Parabruchophagus* and all five known species (*P. kazakhstanicus* Zerova, *P. nikolskaji* (Zerova), *P. rasnitsyni* Zerova, *P. saxatilis* (Zerova), and *P. tauricus* (Zerova)) are transferred back to *Bruchophagus* **syn. nov.**. The species initially described as *Nikanoria* (Nikol'skaya) due to their metallic reflections (although sometimes very slight) are found to be the sister group of the *gibbus* species group. The synonymy of *Nikanoria* with *Bruchophagus*, proposed by Lotfalizadeh *et al.* (2007), and challenged by Zerova, who described additional species in *Nikanoria*, is confirmed here. Finally, *Exeurytoma* Burks, which includes species with a sclerotized body and long, extended ovipositor sheaths, branches within the *gibbus* species group in accordance with the biology of the involved taxa, in which larvae develop within leguminous pods. *Exeurytoma*, which includes three known species (*E. anatolica* Cam, *E. caraganae* Burks, and *E. kebanensis* Doganlar), is therefore synonymized with *Bruchophagus* **syn. nov.**.

**Clade *Eurytoma s.s.***

The distance separating the nodes supporting *Eurytoma*_sp11_USNMENT01525819 (which emerged from a vespid nest in West Africa) + *Masneroma angulifera* and *Eurytoma sensu stricto* is quite reduced, suggesting that *Masneroma* might just be an outstanding form of *Eurytoma*. *Eurytoma* s.s. largely conforms to the species groups defined by Lotfalizadeh et al. (2007), namely the *robusta*, *compressa*, *amygdali*, *morio*, *fumipennis*, and *rosae/arbortani* + *appendigaster* groups (with the backbone poorly resolved). The *amygdali* and *fumipennis* groups include phytophagous species, respectively infesting fruit of Rosaceae and *Euphorbia*. The other groups are assumed to be entomophagous, but they all develop at the expense of concealed hosts; thus, the trophic relationships are often not firmly established. *Eurytoma s.s.* is primarily distributed in the Holarctic region, though species of the *robusta* species group are commonly found in Sub-Saharan Africa.

**References**

Bouček, Z. (1988) Australasian Chalcidoidea (Hymenoptera). A biosystematic revision of genera of fourteen families, with a reclassification of species. 832 pp., CAB International, Wallingford, UK.

Lotfalizadeh, H., Delvare, G., Cruaud, A. & Rasplus J.-Y. (2024) Morphological phylogeny and revision of *Sycophila* and *Ficomila* (Hymenoptera: Chalcidoidea, Eurytomidae) associated with Afrotropical fig trees (Moraceae, *Ficus*). *Zootaxa*, 5401(1): 1–190. https://doi.org/10.11646/zootaxa.5401.1.1

Lotfalizadeh, H., Delvare, G., & Rasplus, J.-Y. (2007). Phylogenetic analysis of Eurytominae (Chalcidoidea: Eurytomidae) based on morphological characters. *Zoological Journal of the Linnean Society*, *151*(3), 441-510. https://doi.org/https://doi.org/10.1111/j.1096-3642.2007.00308.x

Spinola, M. (1811) Essai d'une nouvelle classification générale des Diplolépaires. *Annales du Muséum National d'Histoire Naturelle, Paris*, 17, 38–152.

Yrgu, A. & Delvare, G. (2019) First report of *Paradecatoma bannensis* Masi (Hymenoptera, Eurytomidae as seed parasite of *Cordia africana*. *Phytoparasitica*, 47: 647–657. https://doi.org/10.1007/s12600-019-00763-w

Zerova, M.D. (2010) Palaearctic species of the genus *Eurytoma* (Hymenoptera, Chalcidoidea, Eurytomidae): morphological and biological peculiarities, trophical associations and key to determination. *Vestnik Zoologii*, Supplement 24, 1–203. (in Russian)

Zerova, M.D. (2011) A new status of the subgenus *Parabruchophagus* Zerova, 1992 (Hymenoptera: Eurytomidae) and its composition. *Russian Entomological Journal*, 20(3): 345–350.
